# Host exonuclease SbcB and a phage-encoded SSB-like protein control activation of the DRT10 reverse transcriptase defense system

**DOI:** 10.64898/2026.04.29.721553

**Authors:** Esperanza Sánchez-Nieto, Max E. Wilkinson, Vicenta Millán, Celestine Wenardy, Daniel Cabrerizo, Francisco Martínez-Abarca, Nicolás Toro

## Abstract

Defense-associated reverse transcriptases (DRTs) employ diverse mechanisms of cDNA synthesis to protect bacteria against phage infection, yet their full diversity and regulatory logic remain poorly understood. Here we provide a mechanistic characterization of UG17 (DRT10), a class-2 system within the UG/Abi reverse transcriptase lineage, classifying it into three phylogenetically and architecturally distinct subtypes with subtype-specific ncRNAs and experimentally validating its role in phage defense. DRT10 operates as a tripartite module, comprising a structured ncRNA, a SLATT effector, and an RT that catalyzes processive synthesis of DNA containing 7 nt tandem repeats. The tandem-repeat cDNA intermediate accumulates as both first- and second-strand species. SbcB suppresses DRT10-mediated toxicity and appears to selectively degrade the second strand under basal conditions, while phage infection correlates with enhanced first-strand accumulation. A phage-encoded protein with predicted SSB-like architecture and a conserved C-terminal tip motif is required for efficient DRT10 activation during infection. Together, these findings establish DRT10 as a surveillance system whose activation threshold is jointly controlled by constitutive cDNA synthesis, host exonuclease activity, and a phage-encoded SSB-like trigger that perturbs host ssDNA metabolism. We propose a model in which accumulation of DRT10-derived cDNA triggers activation of the SLATT transmembrane effector to initiate immune defense.

## Introduction

Reverse transcriptases (RTs), enzymes that convert RNA into complementary DNA, were long considered hallmarks of retroviruses, first discovered in RNA tumor viruses in 1970 (Baltimore, 1970; Temin & Mizutani, 1970). The subsequent identification of RTs in bacteria, initially as core components of retrons (Yee *et al*., 1984), revealed that reverse transcription is a far more ancient and widespread process than previously appreciated.

In bacteria, RTs are distributed across multiple functional classes, including group II introns, retrons, diversity-generating retroelements (DGRs), group II-like (G2L) elements, RTs integrated into CRISPR-Cas systems and Abi-like modules, and those classified into unknown groups (UGs) (Simon & Zimmerly, 2008; Toro & Nisa-Martínez, 2014; Zimmerly & Wu, 2015; Toro *et al*., 2019; González-Delgado *et al*., 2021). A growing body of evidence has revealed that many of these prokaryotic RTs, including several UG families, participate in antiviral defense, establishing reverse transcription as an unexpected but widespread component of bacterial immunity (Gao *et al*., 2020; Millman *et al*., 2020; González-Delgado *et al*., 2021; Mestre *et al*., 2022).

UG/Abi systems constitute a major lineage of bacterial defense modules collectively referred to as DRT (defense-associated RT) systems (Gao *et al*., 2020; Mestre *et al*., 2022). These RTs have been classified into three classes encompassing 42 groups, including Abi-A, Abi-K, Abi-P2, and 37 UG families (Mestre et al., 2022). Notably, many of these systems are genetically linked to additional genes or domains encoding putative effector components, and several class 2 UG systems are associated with predicted non-coding RNAs (ncRNAs) (Mestre *et al*., 2022), suggesting that at least a subset of DRT systems operate as multicomponent defense modules rather than standalone enzymes.

To date, nine UG systems have been experimentally validated as antiviral defense systems (DRT1–DRT9), spanning a broad phylogenetic and mechanistic diversity (Gao *et al*., 2020; Mestre *et al*., 2022). Nevertheless, the molecular mechanisms underlying most DRT systems remain poorly understood, and the full functional and mechanistic repertoire of prokaryotic RT-based immunity has yet to be defined.

Structural and mechanistic studies have begun to illuminate how DRT systems operate. The class-1 system DRT6 (UG12) (Mestre *et al*., 2022) and phylogenetically related AbiA, AbiK and Abi-P2 systems consist solely of an RT that assembles into functional hexamers (DRT6, AbiK), dimers (AbiA) and trimers (Abi-P2) (Figiel *et al*., 2022; Gapińska *et al*., 2024; Wang *et al*., 2025). By contrast, several class-2 systems require an essential ncRNA for complex assembly, enzymatic activity, and anti-phage function. In DRT2, the ncRNA serves as the RT template, driving rolling-circle reverse transcription to produce repetitive cDNA molecules that ultimately trigger growth arrest via expression of a toxic open reading frame (Tang *et al*., 2024; Wilkinson *et al*., 2024). In DRT9, long poly-A-rich cDNA molecules likely sequester a phage SSB protein, interfering with phage replication (Han *et al*., 2025; Song *et al*., 2025; Tang *et al*., 2025a). Together, these examples reveal that RT-based defense systems have evolved mechanistically distinct, nucleic-acid-driven strategies, a diversity that likely extends to the many DRT systems that remain uncharacterized.

Among these, the class-2 system UG17 is consistently associated with genes encoding SLATT (SMODS and SLOG-associating 2TM effector) domain proteins, predicted pore-forming toxins proposed to export RT-derived DNA intermediates (Burroughs *et al*., 2015). However, the broader diversity of UG17 systems, their potential role in phage defense, the mechanisms governing cDNA synthesis, and the functional role of the SLATT effector remain unexplored.

Here, we show that UG17 (DRT10) systems function in phage defense and comprise three distinct subtypes defined by RT phylogeny, gene architecture, taxonomic distribution, and ncRNA sequence and structural features. All three subtypes encode tripartite modules composed of a SLATT effector, an RT, and a subtype-specific ncRNA, all of which are required for antiviral activity in the characterized model system. We demonstrate that productive RT-ncRNA interaction, rather than ncRNA stability alone, is a key determinant of system activation, and that the RT catalyzes processive synthesis of tandem-repeat ssDNA through iterative strand repositioning governed by conserved ACU motifs in the ncRNA template. Furthermore, we identify SbcB as a host factor that modulates the cDNA intermediate, and a phage-encoded SSB-like protein as the trigger of the immune response, uncovering previously unrecognized regulatory layers in RT-based immunity.

## Materials and methods

### Bacterial strains, phages, and growth conditions

*Escherichia coli* K-12 MG1655 ΔRM (ATCC 25404) and the BASEL phage collection were kindly provided by Maffei et al. (2021). *E. coli* K-12 MG1655 ΔRM was used as the host for phages from the BASEL collection. *E. coli* K-12 MG1655 (DSM 18039) was obtained from the DSMZ-German Collection of Microorganisms and Cell Cultures GmbH and served as the host for phages T2 (ATCC 11303-B2) and T5 (ATCC 11303-B5), which were acquired from the American Type Culture Collection (ATCC). Strains from the Keio collection (Baba *et al*., 2006) were used as the source for wild-type *E. coli* K-12 BW25113 (OEC5042) and its derivatives Δ*sbcB* (OEC4987-200827348), Δ*recJ* (OEC4987-200828065), and Δ*xseA* (OEC4987-200827798). All *E. coli* strains were grown in Luria-Bertani (LB) medium or on LB agar plates at 37 °C. When required, media were supplemented with ampicillin (Ap, 200 µg mL⁻¹), chloramphenicol (Cm, 50 µg mL⁻¹), and/or tetracycline (Tc, 10 µg mL⁻¹) for plasmid maintenance.

### DNA oligonucleotides

DNA oligonucleotides used for conventional PCR, inverse PCR, blot hybridization, and primer extension were synthesized by Metabion or Integrated DNA Technologies (IDT) and are listed in **Table S1**.

### Plasmid constructions

Plasmids used in this study are listed in **Table S2**. All plasmid constructs were verified by DNA sequencing.

The full-length wild-type UG17-A system from *Escherichia* sp. (E3659) (NZ_VATQ01000016.1) was custom synthesized by GenScript Biotech and subcloned into pUC57mini (p57m), generating p57m_UG17_WT (Ap^R^). SLATT, RT, and ncRNA mutant derivatives were provided by GenScript or generated by subcloning into the same vector (**Table S2**).

RT-UG17 was tagged at both the N- and C-termini with a Flag epitope. The N-terminal Flag fusion (p57m_UG17-WT_Flag-RT) was synthesized by GenScript Biotech. The C-terminal Flag fusion (p57m_UG17-WT_RT-Flag; **Table S2**) was generated by PCR amplification of the Flag sequence and inverse PCR of the p57m_UG17_WT backbone (primers listed in **Table S1**), followed by assembly using the NEBuilder® HiFi DNA Assembly Master Mix (New England Biolabs). RT-UG17 was also tagged at the C-terminus with a 6×His epitope. The construct p57m_UG17-WT_RT-6×His (**Table S2**) was generated by inverse PCR of p57m_UG17_WT using primers introducing the 6×His tag (**Table S1**), followed by XhoI digestion and self-ligation. The 6×His-tagged RT fragment was subsequently transferred into mutant backgrounds (RT-YVAA, ncRNA-SL1-1, ncRNA-SL1-3, and ncRNA-SL1-5) by restriction-based subcloning, generating the corresponding tagged derivatives (**Table S2**).

To generate pBAD30_UG17_WT (**Table S2**), the UG17 locus was subcloned into the pBAD30 vector (Ap^R^) under the control of the arabinose-inducible *araBAD* promoter, regulated by *araC* and repressed by glucose. An intermediate construct (pBAD30_UG17_ncRNA-Δ92; **Table S2**) was generated by restriction-based cloning of the UG17 locus lacking its native promoter. The full-length wild-type UG17 sequence was subsequently amplified by PCR (primers listed in **Table S1**) and inserted into the pBAD30 backbone by restriction digestion and ligation, yielding the final pBAD30_UG17_WT construct.

RT was also tagged at the C-terminus with a 6×His epitope in the pBAD30-based constructs. The 6×His-tagged RT fragment was transferred from the corresponding p57m construct into the pBAD30 backbone by restriction-based subcloning, generating pBAD30_UG17-WT_RT-6×His (**Table S2**). The same strategy was used to introduce the 6×His tag into the SLATT-Δ139 and RT-YVAA mutant backgrounds, yielding the respective tagged derivatives (**Table S2**).

To express Bas52_0087 variants, the wild-type, D252V, and Y351C alleles were amplified from wild-type or mutant phage genomic DNA and cloned into the pBAD33 vector (Cm^R^) under the control of the arabinose-inducible *araBAD* promoter, generating pBAD33_0087-WT, pBAD33_0087-D252V, and pBAD33_0087-Y351C (**Table S2**) by restriction digestion and ligation using primers listed in **Table S1**.

To investigate the role of host factors, the *sbcB* gene encoding ExoI was amplified from *E. coli* K-12 BW25113 genomic DNA and cloned into the pBB3 vector (Tc^R^), generating pBB3_SbcB (**Table S2**). The SbcB15 (A183V) allele was generated by PCR-based mutagenesis and cloned into the same vector, yielding pBB3_SbcB15 (**Table S2**) by restriction digestion and ligation using primers listed in **Table S1**.

For ncRNA complementation, the wild-type ncRNA locus (including its native promoter) was amplified from p57m_UG17_WT and cloned into pBB3, generating pBB3_ESN2 (**Table S2**) by restriction digestion and ligation using primers listed in **Table S1**.

### Bacterial genomic DNA extraction

Bacterial genomic DNA was extracted from 5 mL *E. coli* cultures (OD_600_ = 0.5–0.6) using the REAL Genomic DNA Bacteria kit (REAL Laboratory), according to the manufacturer’s instructions, except that the final DNA pellet was resuspended in 50 µL of Milli-Q® water instead of 200 µL of hydration solution recommended by the manufacturer to obtain a more concentrated preparation. DNA was quantified using a Qubit® 2.0 fluorometer (Invitrogen), and samples were stored at –20 °C until use.

### Phage genomic DNA extraction

Phage genomic DNA was extracted from 1 mL of high-titer lysates (≥10⁹ plaque-forming units (PFU) mL⁻¹), or from 1.5 mL for lower-titer stocks. Lysates were mixed with 0.2 volumes of M1 buffer (3 M NaCl and 30% polyethylene glycol 6000), vortexed briefly, incubated on ice for 1 h, and centrifuged at 13,000 rpm for 10 min at room temperature to pellet phage particles. Pellets were resuspended in 200 µL of M2 buffer (500 mM guanidine-HCl, 10 mM MOPS and 1% Triton X-100; pH 6.5) and incubated for 10 min at 80 °C to lyse phage particles, then cooled to room temperature. Lysates were neutralized with 200 µL of MX3 buffer instead of 350 µL recommended by the manufacturer and purified by centrifugation using mi-Plasmid Miniprep columns (Metabion), according to the manufacturer’s instructions. DNA was eluted in 30 µL of Milli-Q® water instead of 50 µL of elution buffer recommended by the manufacturer to obtain a more concentrated preparation. DNA was quantified using a Qubit® 2.0 fluorometer (Invitrogen), and samples were stored at −20 °C until use.

### Total nucleic acids extraction (DNA and RNA)

Total nucleic acids (DNA and RNA) were extracted from bacterial pellets obtained from 10–60 mL of *E. coli* cultures (OD_600_ = 0.5–0.6). Pellets were resuspended in 300 µL of pre-warmed lysis solution (4 mM Na_2_EDTA·2H_2_O, 130 µg proteinase K (Sigma-Aldrich) and 1.4% SDS) and incubated for 10 min at 65 °C. Proteins were precipitated by adding 150 µL of 5 M NaCl, vortexing briefly, and incubating on ice for 10 min. Samples were centrifuged at 13,200 rpm for 15 min at 4 °C. Nucleic acids were precipitated from the supernatants by incubation for 1 h at –80 °C with 1 mL of absolute ethanol pre-chilled to –20 °C, pelleted by centrifugation at 13,200 rpm for 30 min at 4 °C, and resuspended in 30 µL of Milli-Q® water. Samples were sequentially extracted with phenol:chloroform:isoamyl alcohol (25:24:1; pH 8.0; VWR Life Science) and chloroform:isoamyl alcohol (24:1), followed by precipitation with absolute ethanol pre-chilled to –20 °C in the presence of 3 M sodium acetate (pH 5.2). Pellets were washed with 70% pre-chilled ethanol, air-dried, and resuspended in in 20 µL of Milli-Q® water (Cabanes *et al*., 2000; García-Tomsig *et al*., 2023). RNA integrity and relative abundance were assessed by denaturing agarose gel electrophoresis (1.4% agarose in 1× MOPS buffer containing 0.05% formaldehyde). Gels were run at 100 V and visualized using a GelDoc™ Go Imaging System (Bio-Rad). Samples were stored at –80 °C until use in blot hybridization.

### Total nucleic acid blot hybridization

Total nucleic acid samples (10–20 µg) were denatured in loading buffer (0.3% bromophenol blue, 0.3% xylene cyanol, 10 mM EDTA and 97.5% deionized formamide, adjusted to pH 7.5) for 5 min at 95 °C, cooled on ice, and resolved on 6% denaturing polyacrylamide (acrylamide:bisacrylamide 29:1) gels containing 8 M urea in 1× TBE for 90 min at 20 W. Following electrophoresis, nucleic acids were transferred to positively charged nylon membranes (Roche Diagnostics) by electroblotting. Radiolabeled GeneRuler® DNA molecular weight marker (Thermo Fisher Scientific) was included in each gel. Hybridizations were carried out overnight at 42 °C in phosphate/SDS buffer (10 mM EDTA, 0.5 M Na2HPO4/NaH2PO4 and 7% SDS, adjusted to pH 7.2) using 50 pmol of 5’-end [γ-32P] ATP-labeled oligonucleotide probes generated with T4 polynucleotide kinase (New England Biolabs). Probes used to detect the UG17 ncRNA and *E. coli* 5S rRNA (loading control) are listed in **Table S1**. Membranes were washed by rinsing once with 2× SSC/0.1% SDS at room temperature, followed by three consecutive washes of 15 min each at 42 °C with 2× SSC/0.1% SDS, 1× SSC/0.1% SDS, and 0.1× SSC/0.1% SDS, and exposed to BAS-MP 2040 phosphor imaging screens (Fujifilm). Signals were visualized using a Personal Molecular Imager® FX scanner (Bio-Rad), and images were processed with Quantity One® software (del Val *et al*., 2007; García Tomsig *et al*., 2023).

### Southern blot hybridization

Bacterial cultures either infected with phage or mock-treated were grown from single colonies in LB medium supplemented with appropriate antibiotics at 37 °C to OD600 = 0.5–0.6. For infection experiments, cultures were divided and infected with phage at a multiplicity of infection (MOI) of 5 or mock-treated with LB medium, followed by incubation for 60 min at 37 °C with shaking. Cells were collected by centrifugation at 6,000 rpm for 5 min at 4 °C and pellets stored at –80 °C prior to bacterial genomic DNA extraction as described above. Samples (1–2 µg) were resolved on 0.8% non-denaturing agarose gels in 1× TAE (10× buffer composition: 400 mM Tris, 0.1142% (v/v) glacial acetic acid and 20 mM EDTA) and run overnight at 7 V. Gels were stained with GelRed® (Biotium), visualized under UV light using the GelDoc™ Go Gel Imaging System (Bio-Rad), and then exposed to UV light for 30 min to partially fragment the DNA. Samples were transferred to positively charged nylon membranes (Roche Diagnostics), previously wetted with deionized water and equilibrated for 5–10 min with 20× SSC solution, for 1–2 h in alkaline transfer buffer (1 M NaOH) using the VacuGene XL vacuum blotting system (Pharmacia). Membranes were washed for 10 min with 2× SSC solution with gentle agitation. DNA was immobilized by vacuum heat fixation (70 cm Hg, 120 °C, 35 min). Hybridizations were performed with 5’-end [γ-32P] ATP-labeled oligonucleotide probes as described above. Probes targeting poly-dA products (5’-TTTTTTTTTTTTTTTTTTTTTTTTTTTTTTTTTTTTTTTT-3’), the UG17 first-strand cDNA (5’-TATTACTTATTACTTATTACTTATTACT-3’), and the UG17 second-strand cDNA (5’-AGTAATAAGTAATAAGTAATAAGTAATA-3’) are listed in **Table S1**.

Hybridization and washing were performed at 42 °C, except for the UG17 cDNA probes, which were hybridized and washed at 37 °C due to their lower melting temperature.

### Dot blot hybridization

Bacterial genomic DNA was prepared as described for Southern blot experiments. Samples (1–2 µg) were diluted to a final volume of 40 µL, and serial two-fold dilutions were prepared. Positively charged nylon membranes (Roche Diagnostics) were placed in a Bio-Dot® Microfiltration System (Bio-Rad) and each well was pre-wetted with 20 µL of Milli-Q® water. Aliquots (20 µL) of each diluted sample were applied per well under gentle vacuum. Membranes were air-dried and DNA was immobilized by vacuum heat fixation (70 cm Hg, 120 °C, 35 min). Hybridizations were performed with 5’-end [γ-32P] ATP-labeled oligonucleotide probes as described above. Probes targeting poly-dA products (5’-TTTTTTTTTTTTTTTTTTTTTTTTTTTTTTTTTTTTTTTT-3’), the UG17 first-strand cDNA (5’-TATTACTTATTACTTATTACTTATTACT-3’), and the UG17 second-strand cDNA (5’-AGTAATAAGTAATAAGTAATAAGTAATA-3’) are listed in **Table S1**. Hybridization and washing were performed at 42 °C, except for the UG17 cDNA probes, which were hybridized and washed at 37 °C due to their lower melting temperature.

### 5’-end ncRNA mapping by fluorescent primer extension

The 5’-end of the ncRNA was mapped by fluorescent primer extension using total nucleic acid samples. A 6-FAM-labeled oligonucleotide (5’-6-FAM-CCGAAGGTAGTAATAAGTAACGCCTG-3’; listed in **Table S1**) was annealed to 15 µg of total nucleic acids in annealing buffer (400 mM NaCl and 10 mM PIPES; pH 7.5) by heating for 5 min at 85 °C, rapid cooling to 60 °C, and gradual cooling to 45 °C. Extension reactions were performed by adding 1× AMV RT buffer (Roche Diagnostics), 1 mM dNTPs, 60 µg mL⁻¹ actinomycin D (Sigma), 2 units of murine RNase inhibitor (New England Biolabs), and 7 units of AMV RT (Roche Diagnostics), and incubating for 60 min at 44 °C. Reactions were terminated by ethanol precipitation with 3 M sodium acetate (pH 5.2) (Fekete *et al*., 2003; Muñoz-Adelantado *et al*., 2003; Molina-Sánchez *et al*., 2010; Schuster & Bertram, 2014). Extension products were analyzed by capillary electrophoresis at the Genomics Unit of the Instituto de Parasitología y Biomedicina “López Neyra” (Granada, Spain).

### Double-layer agar assays

Double-layer agar assays were performed as described by Kropinski *et al*. (2009). *E. coli* K-12 MG1655 ΔRM strains were grown overnight at 37 °C in LB medium supplemented with the appropriate antibiotics. Bacterial lawns were prepared by mixing 300 µL of stationary-phase culture with 10 mL of molten top agar (LB containing 0.5% agar and 5 mM MgSO_4_ and antibiotics when required), and pouring onto LB bottom agar plates (12 × 12 cm; 1.5% agar, 5 mM MgSO4 and antibiotics when required). Serial 10-fold phage dilutions were prepared in phage buffer (LB with 5 mM MgSO_4_), and 10 µL of each dilution was spotted onto bacterial lawns. Plates were incubated overnight at 37 °C. Plaques were enumerated and phage titers were calculated as PFU mL⁻¹. When individual plaques were too small to be reliably enumerated, the most concentrated dilution showing no plaques was recorded as corresponding to a single PFU.

To assess the effect of SbcB dosage on DRT10-mediated defense, *E. coli* K-12 MG1655 ΔRM strains expressing the wild-type DRT10 system were co-transformed with either the empty pBB3 vector or pBB3_SbcB expressing *sbcB* constitutively. Phage plaque assays were performed as described above using Bas52 phage, and defense activity was quantified as EOP relative to the empty vector control.

### Phage escape mutants isolation and amplification

Phage plaques arising on lawns of *E. coli* K-12 MG1655 ΔRM expressing the DRT10 system were individually picked and resuspended in 500 µL of phage buffer. Suspensions were incubated for 1 h at room temperature with occasional vortexing to release phage particles from the agar plug, clarified by centrifugation at 4,000 rpm for 10 min, and sterilized by filtration (0.22 µm). Serial ten-fold phage dilution plaque assays were performed on the DRT10-expressing strain and on the isogenic ΔRM control strain lacking DRT10. Escape capacity was assessed by determining the efficiency of plating (EOP), defined as the ratio of PFU mL⁻¹ on DRT10-expressing cells relative to PFU mL⁻¹ on the control strain. Phages exhibiting increased EOP compared to the ancestral wild-type phage on the DRT10 background were considered escape mutants. Single plaques from confirmed escape mutants were amplified in 10 mL of an exponentially growing culture of *E. coli* K-12 MG1655 ΔRM expressing the DRT10 system (OD600 = 0.5–0.6). Cultures were incubated overnight at 37 °C with shaking, and lysates were clarified by centrifugation at 6,000 rpm for 10 min and filtration (0.22 µm) prior to downstream analyses.

### Phage escape mutants whole-genome sequencing and analysis

Genomic libraries were prepared for paired-end Illumina sequencing. Reads were mapped to the Bas52 reference genome (GenBank accession: MZ501102) using Geneious Prime (v2024.0.7). Variants were identified by comparative analysis against the ancestral wild-type phage sequenced in parallel. Only mutations present in the escape mutants and absent from the ancestral phage were considered.

### Cell viability assays with phage-trigger

*E. coli* K-12 MG1655 ΔRM strains carrying either the empty p57m vector (control) or the DRT10 system (p57m_UG17_WT), together with the arabinose-inducible plasmid expressing wild-type or mutant Bas52_0087 variants (pBAD33_0087-WT, pBAD33_0087-D252V, or pBAD33_0087-Y351C) were grown overnight in LB supplemented with the appropriate antibiotics and 1% glucose (non-inducing conditions) at 37 °C. Cultures were diluted to OD600 = 0.5–0.6, and serial 10-fold dilutions were prepared in LB. Aliquots (10 µL) of each dilution were spotted onto LB agar plates containing the appropriate antibiotics and either 1% glucose (no induction) or 1% arabinose (induction). Plates were incubated overnight at 37 °C to assess cell viability.

### Cell viability assays with host-gene backgrounds

To assess the effect of host exonucleases on DRT10-associated toxicity, *E. coli* K-12 BW25113 and its isogenic Δ*sbcB* derivative were transformed with plasmids expressing the wild-type DRT10 system and its SLATT, RT, and ncRNA mutant derivatives. Following transformation, cells were recovered in 1 mL LB for 1 h at 37 °C, and 10-µL aliquots were plated onto LB agar supplemented with the appropriate antibiotics. Plates were incubated overnight at 37 °C to assess viability based on transformant recovery.

To assess specificity, the same assay was performed in Δ*recJ* and Δ*xseA* strains, which lack additional host ssDNA exonucleases (RecJ and ExoVII, respectively). BW25113, Δ*sbcB*, Δ*recJ*, and Δ*xseA* strains were transformed with either the wild-type DRT10 system or the RT-inactive mutant (RT-YVAA), and viability was assessed as described above.

To assess the effect of SbcB catalytic activity on DRT10-associated toxicity, Δ*sbcB* strains expressing the wild-type DRT10 system were co-transformed with plasmids expressing either wild-type *sbcB* (pBB3_SbcB) or the catalytically impaired *sbcB15* allele (pBB3_SbcB15). The same constructs were also introduced into wild-type BW25113 to assess potential dominant negative effects of the *sbcB15* allele. Viability was assessed as described above.

### Affinity purification of RT-containing complexes

*E. coli* K-12 BW25113 strains harboring pBAD30 constructs expressing C-terminal 6×His-tagged RT or the corresponding untagged control were grown overnight in LB supplemented with the appropriate antibiotics and 1% glucose (non-inducing conditions) at 37 °C. Cultures were diluted to OD600 = 0.1 in 30 mL of LB supplemented with the appropriate antibiotics and 1% glucose and grown at 37 °C until OD600 = 0.6. Cells were centrifuged at 4,000 rpm for 4 min at 4 °C to remove glucose, resuspended in 30 mL of LB supplemented with the appropriate antibiotics and 1% arabinose (induction), and incubated for 30 min at 37 °C. Cells were then harvested by centrifugation at 6,000 rpm for 5 min at 4 °C and stored at −80 °C. For p57m-based constructs, cells were grown without induction and harvested at OD600 = 0.5–0.6 by centrifugation at 6,000 rpm for 5 min at 4 °C.

Cell pellets were resuspended in 5 mL of lysis buffer (10% glycerol, 10 mM imidazole, 500 mM NaCl and 25 mM Tris-HCl; pH 8.0) supplemented with protease inhibitors and lysed by sonication with 4 pulses of 15 s each. Clarified lysates were incubated with 100 µL of Ni-NTA agarose (Qiagen), previously equilibrated according to the manufacturer’s instructions, for 2 h at 4 °C with gentle shaking. The resin was washed four times with 500 µL of wash buffer (10% glycerol, 25 mM imidazole, 300 mM NaCl and 25 mM Tris-HCl; pH 8.0) by centrifugation at 3,000 rpm for 1 min at 4 °C. Bound proteins were eluted by incubation with 100 µL of elution buffer (10% glycerol, 200 mM imidazole, 300 mM NaCl and 25 mM Tris-HCl; pH 8.0) for 10 min at 4 °C with gentle shaking, followed by centrifugation at 3,000 rpm for 30 s at 4 °C. Eluates were stored at −80 °C.

To assess RT–ncRNA association, eluates were treated with 200 µg proteinase K for 30 min at 37 °C to remove proteins. ncRNA was extracted sequentially with one volume of phenol:chloroform:isoamyl alcohol (25:24:1) and one volume of chloroform:isoamyl alcohol (24:1), each followed by centrifugation at 13,200 rpm for 5 min at 4 °C. ncRNA was precipitated by incubation for at least 1 h at −80 °C with 3 volumes of absolute ethanol, 30 µg glycogen (Roche), and 1/10 volume of 3 M sodium acetate (pH 5.2), followed by centrifugation at 13,200 rpm for 30 min at 4 °C. Pellets were washed with 1 mL of 70% pre-chilled ethanol, air-dried, and resuspended in 10 µL of Milli-Q® water until use in blot hybridization.

### Strand-specific cDNA sequencing

For *in vivo* cDNA sequencing, total DNA was isolated from *E. coli* cultures as described above. For in vitro cDNA sequencing, DNA products were extracted from purified DRT10 RNP complexes following incubation with dNTPs, using proteinase K digestion followed by phenol:chloroform:isoamyl alcohol extraction and ethanol precipitation.

Strand- and end-specific DNA sequencing libraries were prepared as described previously (Khan *et al*., 2024; Wilkinson *et al*., 2024). The mixture was incubated with terminal transferase (New England Biolabs) and 4 µM dATP to dA-tail 3’ ends. An anchored primer (5’-GTCTCGTGGGCTCGGAGATGTGTATAAGAGACAGTTTTTTTTTTV-3’) was annealed to the poly-A tails and extended with exo- Klenow polymerase (New England Biolabs). Reaction products were purified using QIAquick PCR cleanup columns (Qiagen) and ligated to pre-annealed adapters (top strand: 5’-TCGTCGGCAGCGTCAGATGTGTATAAGAGACAGT-3’; bottom strand: 5’- /5Phos/CTGTCTCTTATACACATCTGACGCTGCCGACGA/3AmMO/-3’) using Blunt/TA Ligase Master Mix (New England Biolabs). Libraries were PCR-amplified with indexed primers and sequenced on a MiSeq or NovaSeq X Plus in paired-end mode (2 × 150 bp).

### Bioinformatic analysis of sequencing reads

Sequencing reads were trimmed using Cutadapt (Martin, 2011) to remove adapter sequences and low-quality bases, and mapped to the *E. coli* K-12 genome and expression plasmid sequences using Bowtie2 (Langmead & Salzberg, 2012). Unmapped reads were extracted for further analysis. To identify enriched sequence motifs in DRT10-dependent cDNA products, unbiased 7-mer analysis was performed on unmapped reads by breaking each read into overlapping sequential 7-mers and counting their frequency, normalized by total read count. Enrichment values were calculated as the ratio of normalized 7-mer frequencies between RT-active and RT-inactive samples. To characterize the precision of tandem-repeat synthesis, transition-probability analysis was performed on cDNA reads containing the AGTAATAAGT sequence, quantifying the probability of each base transition within the repeat unit.

### Representative UG17 dataset construction

To expand the repertoire of known UG17 RTs, we downloaded the most up-to-date bacterial RT sequences (annotated as ‘reverse transcriptase’) available in the NCBI database as of October 10, 2024, for a total of 107,067 sequences. To reduce redundancy, the sequences were clustered using CD-HIT (Fu *et al*., 2012) with default parameters and the -c 0.90 option, representing a 90% average amino acid identity (AAI) threshold. This clustering yielded 26,799 non-redundant clusters, each of which represented a unique RT variant. Members of the UG17 RT group were identified using hidden Markov model (HMM) search. An HMM profile specific to UG17 RTs was constructed using *hmmbuild* (part of the HMMER 3.3 suite; Eddy, 2011) based on an alignment of RT sequences from a previously curated UG17 dataset (Mestre *et al*., 2022). The *hmmsearch* tool was used to query clustered RT sequences against the UG17 HMM profile. A total of 484 sequences with significant matches (E-value <10⁻^100^) were classified as UG17 RTs. To confirm their classification, these candidate UG17 RT sequences were aligned with members of the previously reported UG17 dataset, and the resulting multiple sequence alignment (MSA) was subjected to phylogenetic analysis. The corresponding complete and whole-genome shotgun (WGS) sequences encoding these RTs were subsequently retrieved from the NCBI database, and their SLATT-associated encoding genes were extracted for further phylogenetic analyses using custom Python scripts.

### Phylogenetic analyses

RT- and SLATT-associated protein sequences from the novel UG17 dataset were aligned using MAFFT (Katoh & Standley, 2013), and phylogenetic trees were constructed with FastTree (Price *et al*., 2010) using the WAG evolutionary model and a discrete gamma model with 20 rate categories. To assess whether the observed subtype-associated clustering reflects deep phylogenetic divergence of the RT core, additional maximum-likelihood phylogenies were inferred with IQ-TREE2 (Minh *et al*., 2020). For IQ-TREE2 analyses, alignments were trimmed using trimAl (Capella-Gutiérrez *et al*., 2009) with a custom Python script to retain only the conserved RT core domain prior to tree inference. For these analyses, the best-fitting amino acid substitution model was selected independently for each alignment using ModelFinder, and branch support was evaluated using ultrafast bootstrap approximation and SH-aLRT tests (1,000 replicates each). In the SLATT phylogenetic tree, sequences of SLATT-associated proteins from some loci belonging to the DRT3 (UG3–UG8) system, classified within the same protein cluster (Cluster 7; Mestre *et al*., 2022), were included as an outgroup. The same phylogenetic approach was applied to classify additional RT–SLATT loci, including the prophage-encoded system from uropathogenic *E. coli* H120 and the PICI-encoded element KpCIUCICRE8, within the UG17 subtype framework. Phylogenetic trees were visualized and annotated using the ggtree package in R (Yu *et al*., 2017).

### ncRNAs structure prediction

Comparative sequence analyses of the intergenic regions for each UG17 subtype suggested the presence of a conserved ncRNA upstream of the SLATT-encoding gene. MLocarna (Will *et al*., 2007) was used to compute a multiple sequence alignment of these hypothetical RNA sequences, which were then evaluated for statistically significant covarying pairs using R-scape at an *E*-value threshold of 0.05 (Rivas *et al*., 2020). The cascade variation/covariation constrained folding algorithm (CaCoFold) method (Rivas, 2020) was used to improve the predicted ncRNA consensus structures. Covariance models were then built using *cmbuild* and *cmcalibrate* from the Infernal suite (Rivas *et al*., 2020), and applied to the complete and whole-genome shotgun sequences carrying the identified UG17 systems in the constructed database using *cmsearch* from the same suite.

Paired RNA-seq reads (SRR4101233) from *Methylotenera mobilis* JLW8, and (SRR5307609) from *Exiguobacterium* sp. ATb1 were downloaded from the NCBI SRA archive and aligned to their respective genomes using Bowtie2 (Langmead & Salzberg, 2012). For *Escherichia* sp. E3659 UG17 system, RNA-seq was performed. *E. coli* K-12 MG1655 ΔRM cells harboring the synthesized E3659 UG17 system (wild-type and RT-inactive) cloned in p57m were grown to an OD_600_ = 0.5–0.6 in LB medium supplemented with the appropriate antibiotics, and RNA-seq was performed as previously reported (Mestre *et al*., 2022).

### Structural and functional prediction of Bas52_0087

The amino acid sequence of Bas52_0087 was submitted to Phold (Bouras *et al*., 2024) for functional annotation, which predicted an SSB-like fold. Three-dimensional structure prediction was performed using AlphaFold2 (Jumper *et al*., 2021). The predicted structure was visualized and analyzed using PyMOL. Escape mutant substitutions D252V and Y351C were mapped onto the predicted structure to identify their spatial localization relative to predicted functional domains.

## Results

### Phylogenetic and genomic analyses reveal three distinct subtypes within the UG17 system

UG17 systems encode a SLATT domain protein of the SLATT-5 family together with a reverse transcriptase (RT) (Burroughs *et al*., 2015). These systems exhibit two distinct gene architectures based on the relative positioning of the SLATT and RT genes. In the predominant SLATT-RT arrangement, the two genes are tightly coupled, with overlapping stop and start codons. In the alternative RT-SLATT arrangement, the RT gene is positioned upstream of the SLATT gene, separated by approximately 200 bp, with putative promoters identifiable upstream of both genes, suggesting partially independent transcriptional regulation. This architectural variability provided an initial basis for exploring the diversity of UG17 systems.

Phylogenetic analysis of 484 UG17 RT sequences revealed three well-supported groups (**Figure 1A**). Two of these groups correspond to the two previously described gene architectures (Mestre *et al*., 2022), while the third, which also follows the SLATT-RT arrangement, is largely restricted to the phylum Bacillota.

**Figure 1.**
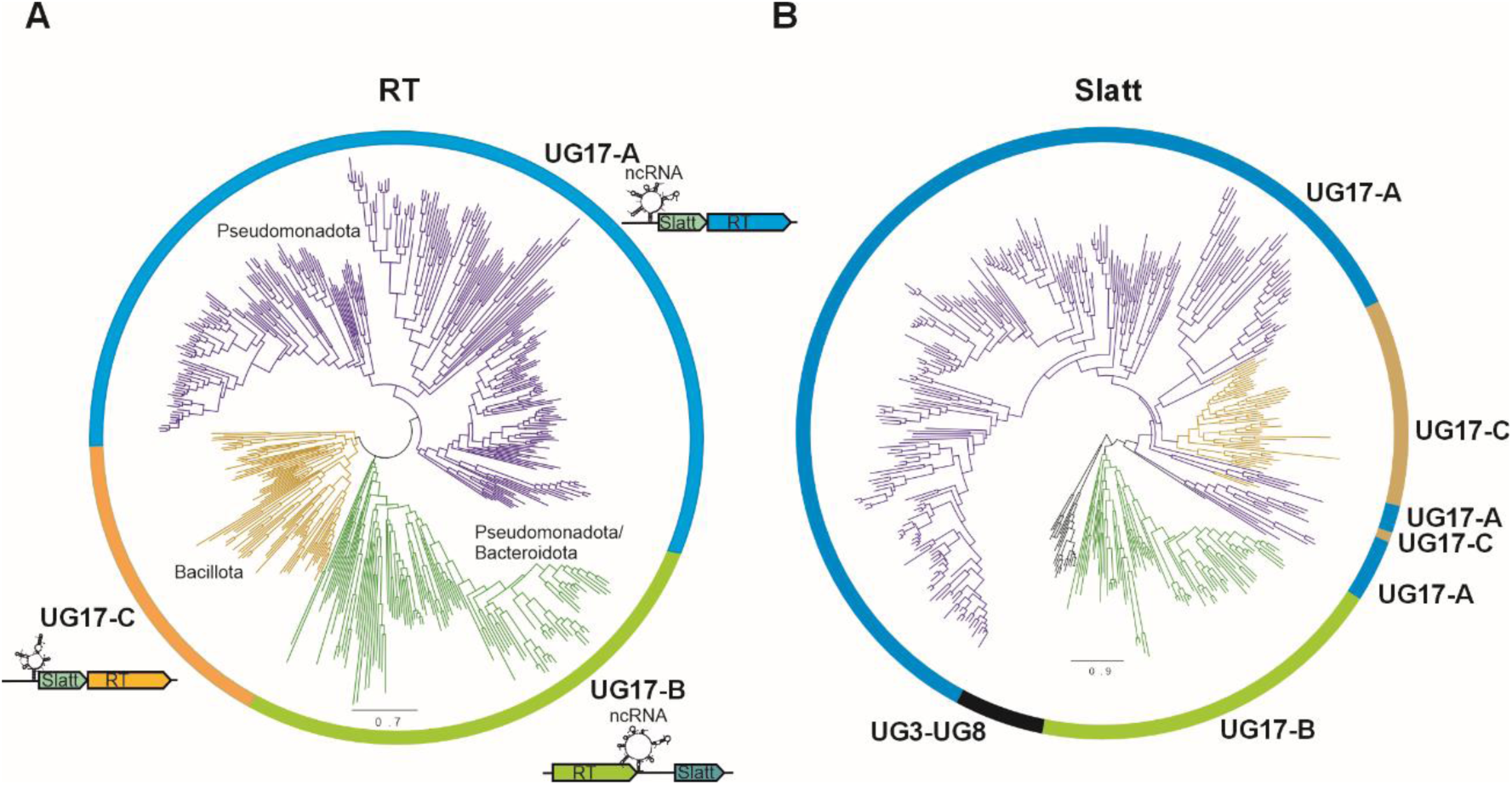
Phylogenetic trees of RT and SLATT protein sequences in the UG17 systems. RT and SLATT sequences were aligned using MAFFT, and the resulting alignments were used as input for FastTree. (**A**) Maximum-likelihood phylogenetic tree of 484 UG17 RT sequences inferred with FastTree. Major sequence clusters are indicated together with their associated organizational architectures. The UG17 subtypes, defined based on RT clustering, domain architecture, taxonomic distribution, and associated ncRNA features, are shown. Predicted ncRNA elements (Figure 2) and the predominant phylum associated with each subtype are also indicated. (**B**) Maximum-likelihood phylogenetic tree of 413 UG17 SLATT protein sequences inferred with FastTree and rooted using 20 divergent SLATT protein sequences from the UG3-UG8 (DRT3) system as an outgroup. Newick tree files for panels A and B are provided as **Supplementary Data Files 1 and 2**, respectively.

Phylogenetic analysis of the associated SLATT proteins revealed a topology that does not mirror the three RT-defined groups (**Figure 1B**), indicating incongruence between the evolutionary histories of the RT and SLATT components. This finding suggests that the two proteins have diversified independently, consistent with the modular architecture of UG17 systems. Phylogenetic analysis restricted to the conserved RT catalytic core did not recover the three subtypes as strictly monophyletic clades (**Figure S1**), suggesting that subtype diversification extends beyond the catalytic domain.

Given that UG loci closely related to UG17 like UG2 and UG3 encode ncRNAs (Mestre et al., 2022), we wondered whether the UG17 subtypes encode associated ncRNAs. Indeed, computational RNA folding of intergenic sequences upstream of the SLATT gene predicted short structured ncRNAs in representatives of all three groups, which were supported by covariance models. However, these models were effective only within individual subtypes (**Figures 2A–C**), reflecting the substantial sequence divergence among UG17-associated ncRNAs.

**Figure 2.**
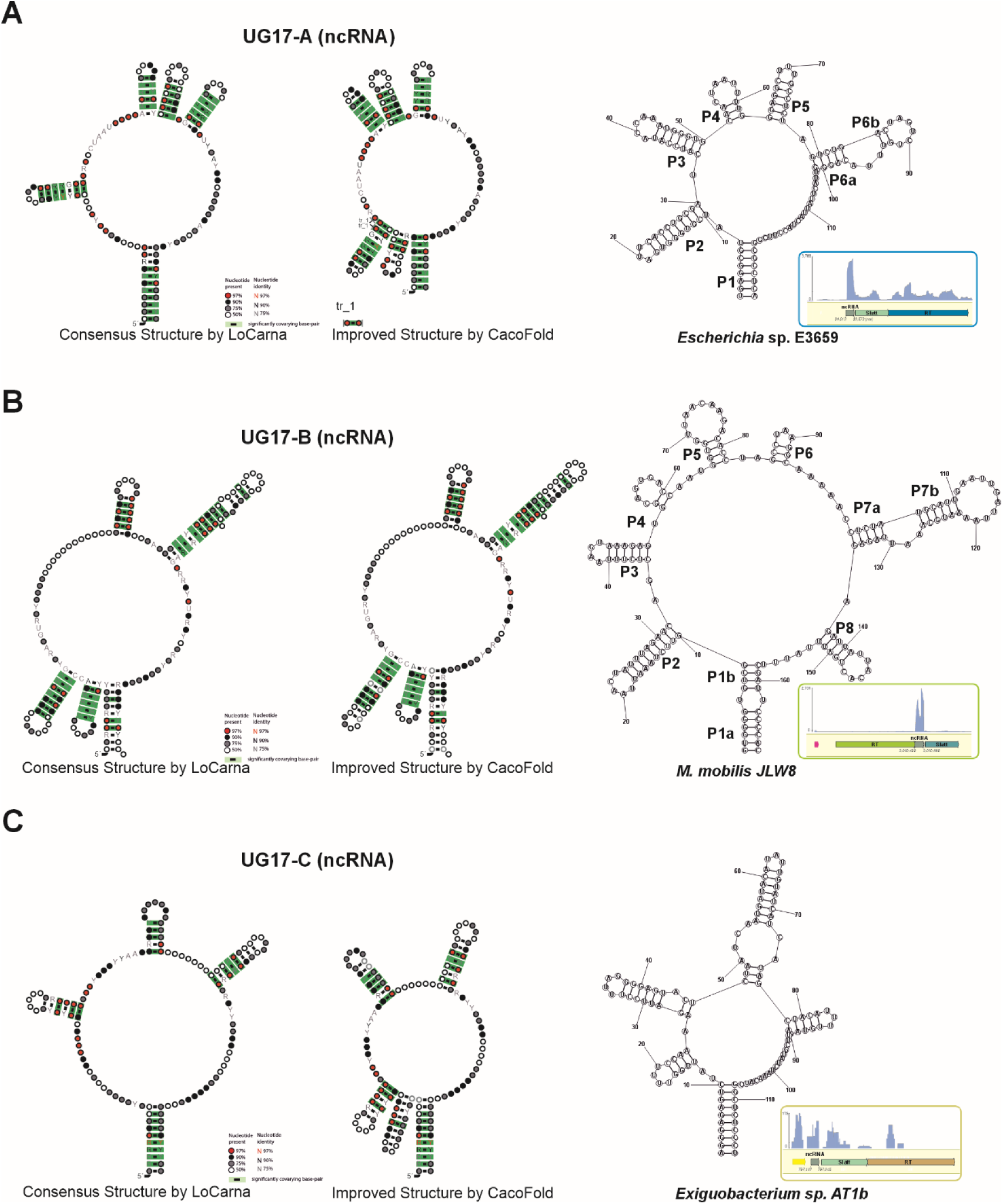
Predicted ncRNA structures of the UG17 subtypes. **A, B, and C.** Predicted ncRNA structures generated by MLocarna and refined using CaCoFold with R-scape are shown. RNA structures of representative UG17 members obtained with RNAstructure (Reuter & Mathews, 2010) are presented together with their RNA-seq alignments, providing evidence for their expression. (**A**) UG17-A subtype: predicted ncRNA structure and RNA-seq alignment for the *Escherichia sp.* E3659 isolate, including the predicted triplet (tr_1). (**B**) UG17-B subtype: predicted ncRNA structure and RNA-seq alignment for the *Methylotenera mobilis* JLW8 strain. (**C**) UG17-C subtype: predicted ncRNA structure and RNA-seq alignment for the *Exiguobacterium sp.* AT1b isolate.

Based on the combined evidence from RT phylogeny, gene architecture, taxonomic distribution, and ncRNA sequence and structural features, we classified UG17 systems into three subtypes: UG17-A, predominantly found in Pseudomonadota; UG17-B, associated with both Pseudomonadota and Bacteroidota; and UG17-C, largely restricted to Bacillota (**Figure 1A**). Together, these analyses establish UG17 as a diverse family of tripartite modules whose RT, SLATT, and ncRNA components have undergone evolutionary diversification.

### A UG17-A model system encodes an independently expressed structured ncRNA stabilized by a conserved basal stem-loop

Alignment of publicly available RNA-seq data to representative UG17-B and UG17-C loci confirmed ncRNA expression upstream of the SLATT gene in both subtypes (**Figures 2A–B**), establishing the ncRNA as a conserved feature across UG17 systems. To characterize the ncRNA component in detail, we focused on a representative UG17-A system from *Escherichia* sp. E3659 as a model, for which we performed RNA-seq analysis alongside structural and functional characterization. This system is phylogenetically related to the UG17-A locus from *Klebsiella* sp. GW_Kp182 (NZ_JBEAFQ010000040.1) in our expanded UG17 dataset. RNA-seq data confirmed ncRNA expression immediately upstream of the SLATT gene (**Figure 2C**), which is followed by the RT gene (**Figure 3A**), an organization consistent with the ncRNA functioning as an integral component of the tripartite UG17-A module.

**Figure 3.**
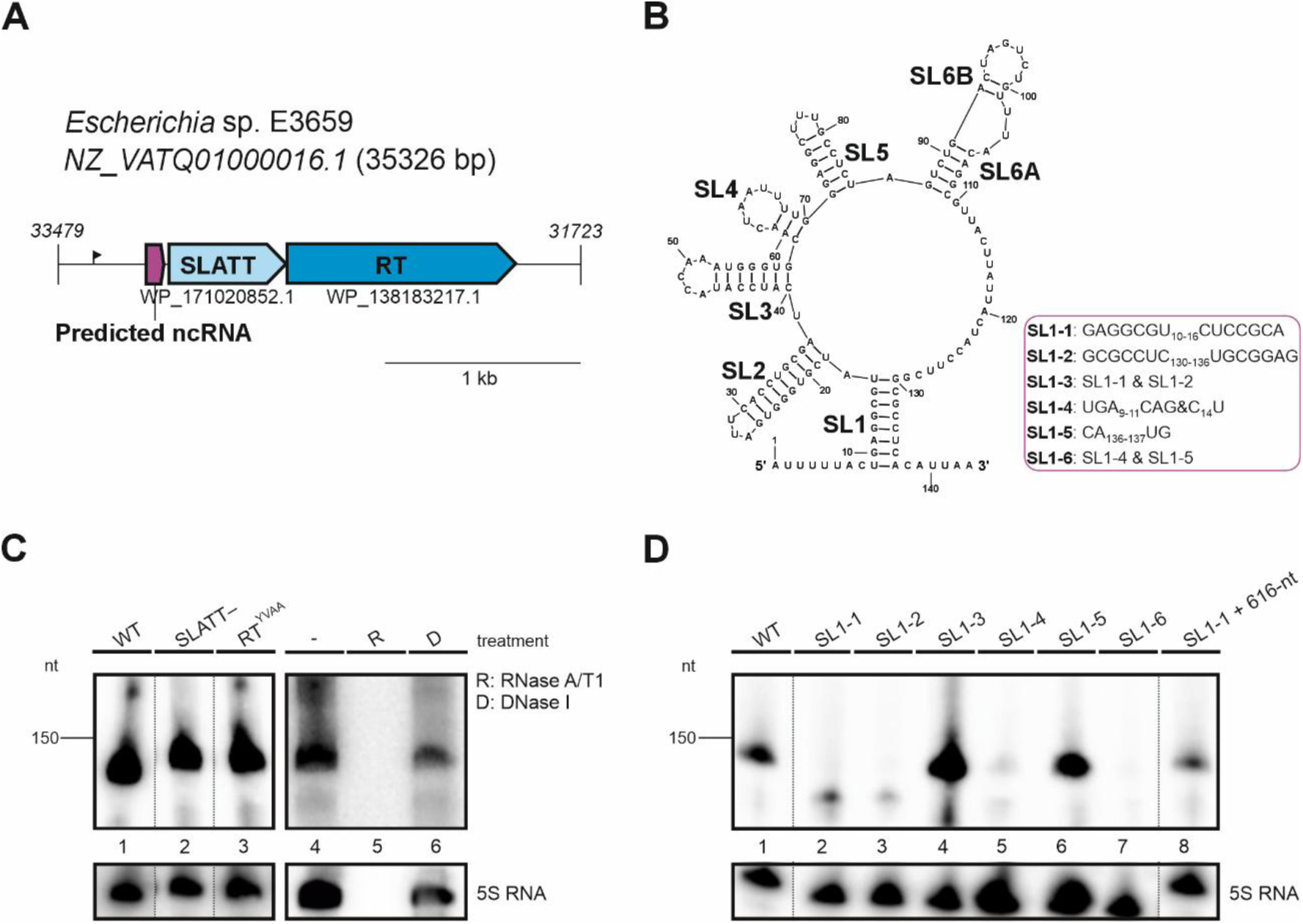
Genomic context, structure, and expression of the UG17-A ncRNA. (**A**) Schematic representation of the UG17-A tripartite system identified in the *Escherichia* sp. E3659 genome (NZ_VATQ01000016.1), comprising an ncRNA (coordinates: 34,014–33,872), a SLATT domain-containing protein (33,857–33,297), and a reverse transcriptase (RT; 33,297–31,960). A black arrow indicates a putative promoter. (**B**) Predicted secondary structure of the 143-nt ncRNA, featuring six stem-loop (SL) structures. Note that the precise boundaries and secondary structure of the ncRNA remain to be established, as additional structural data may refine both the size and the number of stem-loop elements. Mutations introduced in the basal SL1 stem used in different assays are indicated. (**C**) Total nucleic acid blot analysis of *E. coli* K-12 BW25113 cultures expressing the wild-type UG17-A system (WT; lane 1) or the mutant constructs (SLATT− and RT^YVAA^; lanes 2 and 3). Lane 4 shows the untreated control. The ncRNA signal was abolished by RNase A/T1 treatment (lane 5) but persisted following DNase I treatment (lane 6). (**D**) Total nucleic acid blot analysis of *E. coli* K-12 BW25113 cultures expressing the wild-type UG17-A system (WT; lane 1) or the SL1 mutant constructs (SL1-1 to SL1-6; lanes 2−7), and SL1-1 complemented *in trans* with the pBB3_ESN2 plasmid containing the 616-nt ncRNA fragment (616-nt; lane 8). Equal amounts of total nucleic acid were loaded per lane (OD_600_ = 0.5–0.6). Blots were probed with a specific oligonucleotide targeting the predicted ncRNA (5’-GGATGATCGCAGGTGAATCACCCACG-3’). A probe against *E. coli* 5S RNA (5’- TACTCTCGCATGGGGAGACCCCAC-3’) was used as an RNA loading control. The sizes of the 5’-end-labelled GeneRuler DNA Molecular Weight Marker are shown on the left-hand side of each panel.

Secondary structure prediction of the ncRNA revealed six stem-loop structures (SL1–SL6), with SL1 forming the basal stem of the molecule (**Figure 3B**). To experimentally validate the ncRNA and estimate its size, we performed northern blot and primer extension analyses of total nucleic acids, detecting a discrete RNA species of approximately 143-nt (**Figure 3C**) consistent with the predicted structure. However, the precise boundaries and secondary structure of the ncRNA remain to be established, as additional structural data may refine both the size and folding of the molecule. The signal persisted after DNase I treatment but was abolished by RNase A/T1 digestion, confirming the identity of the detected molecule as a bona fide RNA. Notably, ncRNA accumulation was unaffected by mutations in either the SLATT or RT genes, indicating that its expression is independent of the downstream protein-coding components of the UG17-A module.

To assess the contribution of SL1 to ncRNA stability, we introduced mutations disrupting its base pairing (**Figures 3B** and **Figure S2**). Disruption of SL1 base pairing abolished or severely reduced ncRNA accumulation, while compensatory mutations restoring base pairing rescued accumulation to near wild-type levels (**Figure 3D**), demonstrating that SL1 structural integrity is essential for ncRNA stability. Notably, one double mutant that preserved SL1 base pairing nonetheless failed to restore ncRNA accumulation (**Figure 3D**), indicating that specific nucleotide identities within SL1, in addition to its secondary structure, contribute to ncRNA stability.

To further confirm the transcriptional independence of the ncRNA, we expressed it *in trans* under its native promoter in the SL1-1 background. This complementation fully restored ncRNA accumulation (**Figure 3D**), establishing that the ncRNA is expressed as a discrete, independently transcribed component of the UG17-A module.

Having established the ncRNA as a discrete, structurally organized component of the UG17-A module, we next investigated whether the complete tripartite system confers antiviral defense activity.

### All three components of the UG17-A module are required for antiviral defense

To characterize the antiviral activity of our UG17-A model system, we cloned the locus from *Escherichia* sp. E3659 into *E. coli* K-12 MG1655 ΔRM and assessed defense against a panel of 63 phages from the BASEL collection (Maffei *et al*., 2021). The system conferred robust protection against multiple phage subfamilies, including Queuovirinae, Markadamsvirinae, Vequintavirinae, and Stephanstirmvirinae (**Figure 4A**), demonstrating broad-spectrum antiviral activity against diverse tailed phages.

**Figure 4.**
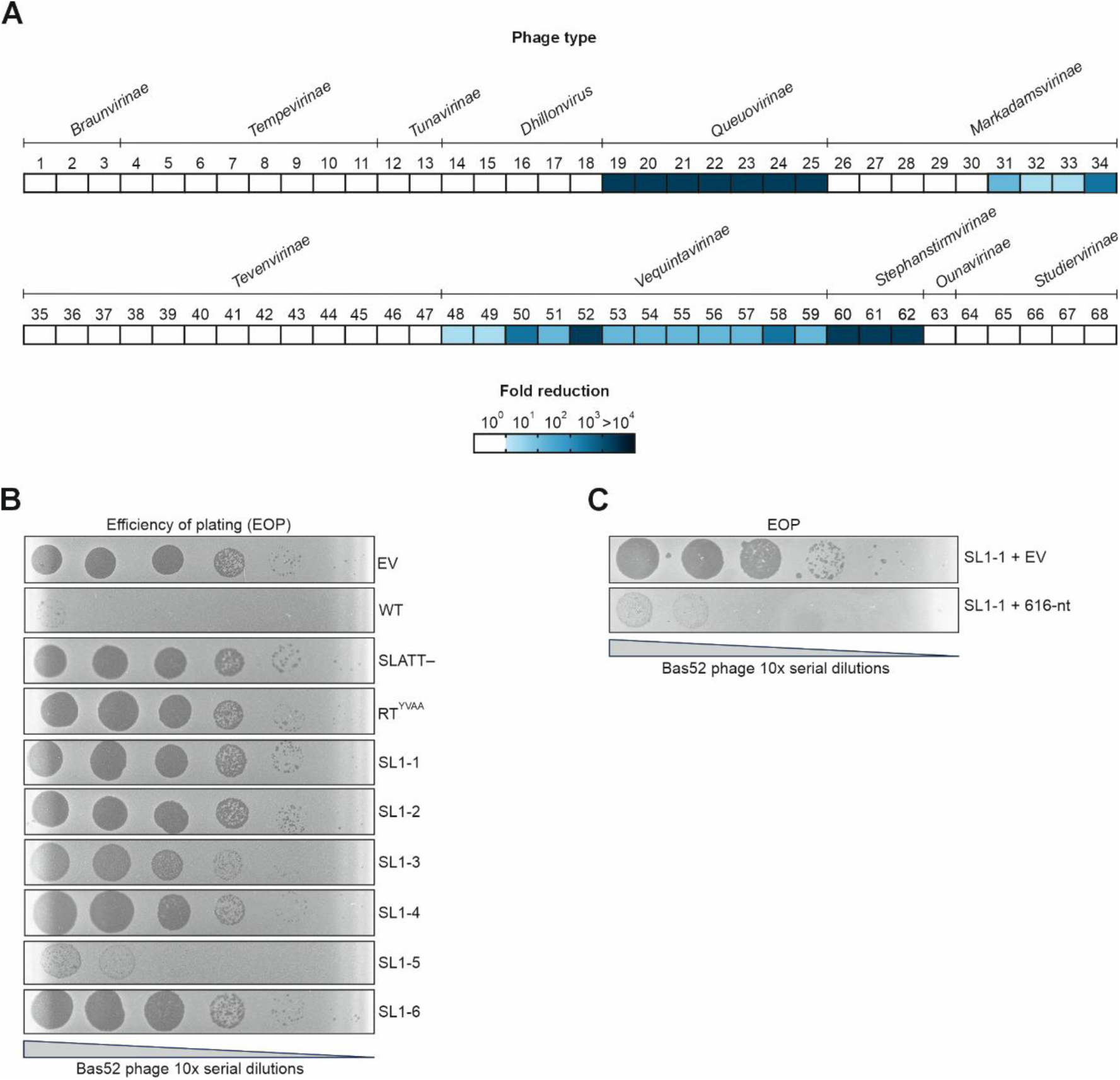
Anti-phage defense activity of the DRT10 system. (**A**) Heat map summarizing the fold-reduction in phage infectivity conferred by the DRT10 system against the 63 phages of the BASEL collection, taxonomically classified according to their subfamilies or genera. Fold reduction was determined by plaque assays using serial 10-fold dilutions, comparing the efficiency of plating (EOP) between *E. coli* K-12 MG1655 ΔRM strains lacking or carrying the DRT10 system. (**B**) Serial 10-fold dilution plaque assays comparing the EOP of phage Bas52 on *E. coli* K-12 MG1655 ΔRM strains expressing the control pUC57m empty vector (EV), the wild-type DRT10 system (WT), or the mutant constructs (SLATT−, RT^YVAA^, and SL1-1 to SL1-6). For clarity, only plaques corresponding to Bas52 are shown. (**C**) Serial 10-fold dilution plaque assays comparing the EOP of phage Bas52 on *E. coli* K-12 MG1655 ΔRM strains expressing the SL1-1 mutant with either the control pBB3 empty vector (EV) or the pBB3_ESN2 plasmid containing the 616-nt ncRNA fragment (616-nt). Data are representative of three independent experiments.

We selected Bas52 as a model phage for subsequent analyses based on its strong and reproducible restriction in the initial screen. Mutation of the SLATT effector, the RT catalytic site, or the ncRNA basal stem all abolished or severely impaired defense against Bas52 (**Figure 4B**), and complementation of the ncRNA mutant in trans fully restored protection (**Figure 4C**), confirming that all three components are essential for antiviral activity. Based on these findings, we refer to this system hereafter as DRT10.

### The host exonuclease SbcB suppresses DRT10-mediated toxicity

A genome-wide essentiality study in uropathogenic *E. coli* identified a genetic interaction between *sbcB* and a prophage-encoded RT–SLATT locus (Rousset *et al*., 2021). SbcB encodes the 3’→5’ ssDNA exonuclease ExoI, and repression of *sbcB* caused a fitness defect in strains expressing this system. Phylogenetic analysis of the RT and SLATT components places this locus within the UG17-B subtype (**Figure S3**), functionally linking it to DRT10 and suggesting that SbcB-encoded ExoI modulates DRT10 activity.

To directly examine the role of SbcB in DRT10 activity, we expressed the DRT10 system in wild-type *E. coli* K-12 BW25113 and its isogenic Δ*sbcB* derivative. While transformants were readily obtained in the wild-type background, no viable transformants were recovered in the Δ*sbcB* strain, indicating that DRT10 activity is toxic in the absence of SbcB. This toxicity required an active tripartite system, as Δ*sbcB* cells tolerated all mutations that inactivated the SLATT effector, the RT catalytic site, or the ncRNA (**Figure 5A**), whereas complementation of the ncRNA mutant in trans with the wild-type ncRNA locus fully restored toxicity in the Δ*sbcB* background (**Figure 5B**).

**Figure 5.**
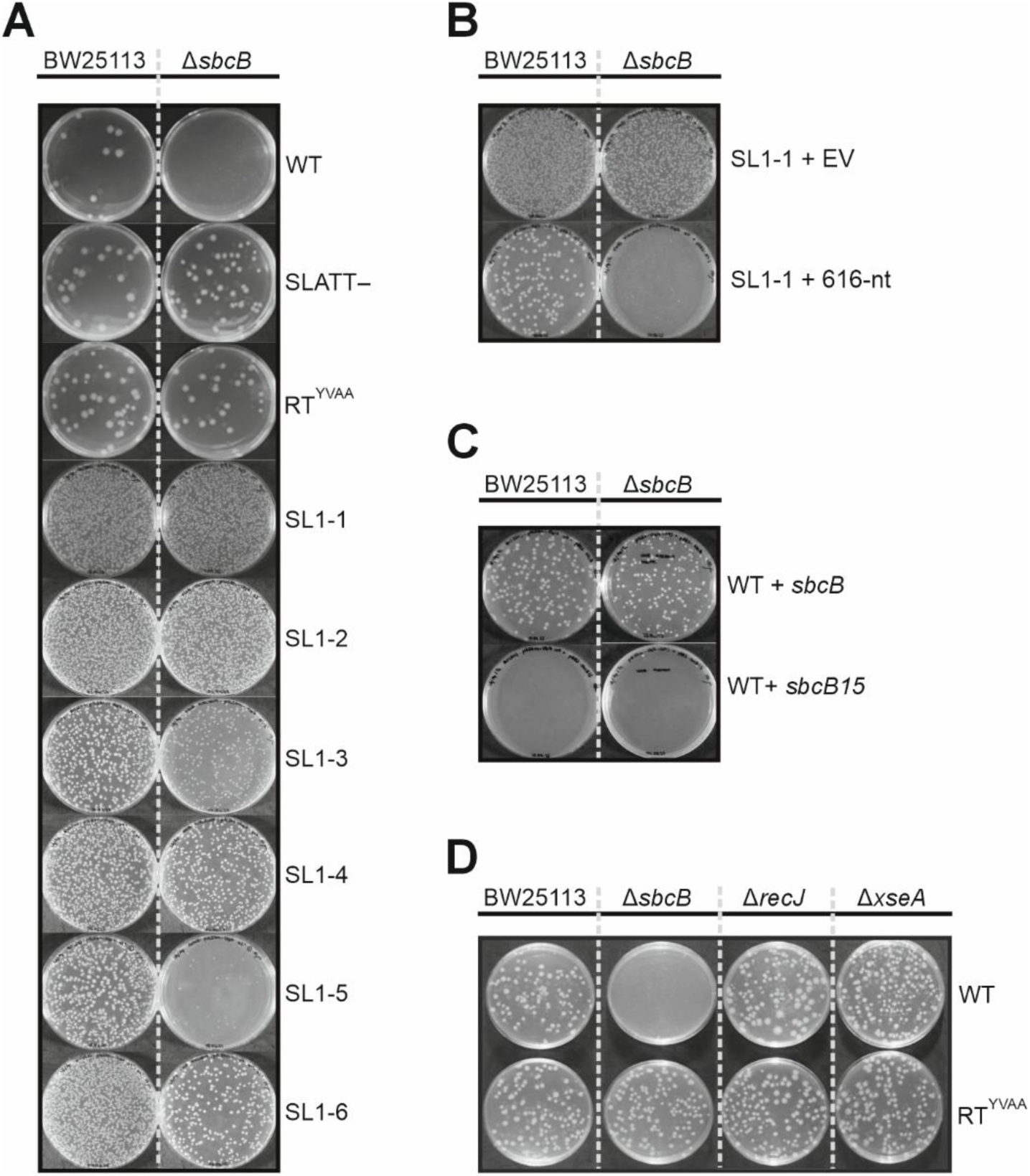
SbcB suppress DRT10-mediated toxicity. (**A**) Transformation assays of *E. coli* K-12 BW25113 and its isogenic Δ*sbcB* derivative with the wild-type DRT10 system (WT), or the mutant constructs (SLATT−, RT^YVAA^, and SL1-1 to SL1-6). (**B**) Co-transformation of *E. coli* K-12 BW25113 and Δ*sbcB* strains with the SL1-1 mutant and either the control pBB3 empty vector (EV) or the pBB3_ESN2 plasmid containing the 616-nt ncRNA fragment (616-nt). (**C**) Co-transformation of *E. coli* K-12 BW25113 and ΔsbcB strains with the wild-type DRT10 system (WT) and either pBB3_SbcB or pBB3_SbcB15 expressing wild-type *sbcB* or the catalytically impaired *sbcB15* variant (A183V), respectively. (**D**) Transformation of *E. coli* K-12 BW25113, Δ*sbcB*, Δ*recJ*, or Δ*xseA* strains with either the wild-type DRT10 system (WT) or the RT-inactive mutant (RT^YVAA^). Viability was assessed by transformant recovery on LB agar plates supplemented with the appropriate antibiotics. Data are representative of three independent experiments.

To confirm that SbcB activity is responsible for suppressing DRT10-mediated toxicity, we complemented the Δ*sbcB* strain with either wild-type *sbcB* or the *sbcB15* allele, which encodes a catalytically impaired ExoI variant (Thoms *et al*., 2008). While wild-type SbcB fully restored viability, the catalytically impaired variant failed to do so. Notably, expression of *sbcB15* also impaired viability in the wild-type BW25113 background (**Figure 5C**), suggesting that the *sbcB15* allele acts in a dominant negative manner over the endogenous SbcB, further supporting the requirement for ExoI catalytic activity in suppressing DRT10-mediated toxicity.

The toxicity phenotype was specific to SbcB, as deletion of other host ssDNA exonucleases, RecJ or ExoVII, had no effect on DRT10-associated lethality (**Figure 5D**). Given that SbcB is a 3’→5’ ssDNA exonuclease, this specificity suggests that DRT10 generates a ssDNA intermediate with a free 3’ end that requires SbcB-mediated degradation for cellular tolerance.

### Productive RT-ncRNA interaction, rather than ncRNA stability, determines DRT10 activity

To assess RT-ncRNA interactions within the DRT10 system, we generated C-terminally 6×His-tagged RT variants that retained partial but detectable defense activity (**Figure S4A–B**), and used these to perform co-immunoprecipitation followed by detection of the ncRNA by blot hybridization (**Figure 6**). The ncRNA was readily detected in pull-downs from wild-type cultures, and this association was unaffected by mutations in the SLATT effector or the RT catalytic site. The SL1-3 double mutant, which preserves ncRNA structure and expression levels, failed to support RT-ncRNA association, consistent with the lack of phage defense in this mutant. This pattern mirrors the differential toxicity observed in the Δ*sbcB* background, establishing that RT-ncRNA association, rather than ncRNA accumulation alone, determines functional system output. Together, these results demonstrate that productive RT-ncRNA interaction is a key determinant of DRT10 activation, and that specific structural features of the ncRNA beyond its stability are required for complex formation with the RT.

**Figure 6.**
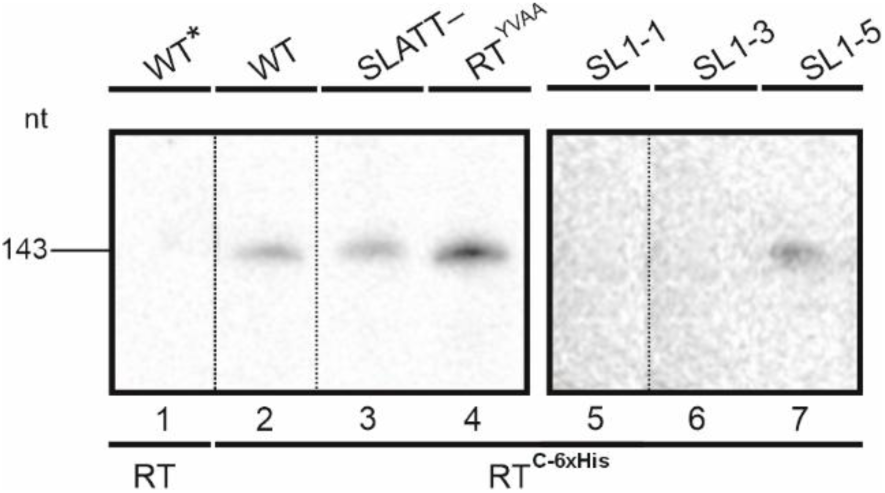
RT-ncRNA association depends on specific ncRNA structural features beyond stability. Co-immunoprecipitation experiments using C-terminally 6×His-tagged RT-UG17 in *E. coli* K-12 BW25113 cultures expressing the control wild-type DRT10 system with untagged RT (WT*; lane 1), the wild-type DRT10 system with the tagged RT (WT; lane 2), or the mutant constructs with the tagged RT (SLATT−, RT^YVAA^, SL1-1, SL1-3, and SL1-5; lanes 3−7). All samples were normalized to the same cell density (OD_600_ = 0.5–0.6). The 143-nt ncRNA bound to the RT was detected by northern blot hybridization using a specific oligonucleotide probe listed in **Table S1**. The estimated size is shown on the left-hand side.

### DRT10 synthesizes 7-mer repetitive DNA

Given that DRT10 requires an ncRNA for antiviral activity, we asked whether this RNA serves as a template for cDNA synthesis. We first tested for poly-dA homopolymers characteristic of DRT9, but no such products were detected in DRT10-expressing strains under phage infection conditions (**Figure S5**), suggesting that DRT10 employs a distinct cDNA synthesis pathway.

To identify the cDNA products generated by DRT10, we purified the DRT10 RNP complex, and sequenced any copurified DNA using strand-specific DNA sequencing. Although no reads mapped to the ncRNA sequence, analysis of unmapped reads revealed that approximately 20% contained tandem copies of a 7-nt AGTAATA motif arranged in extended repeat tracts (**Figure 7A**).

**Figure 7.**
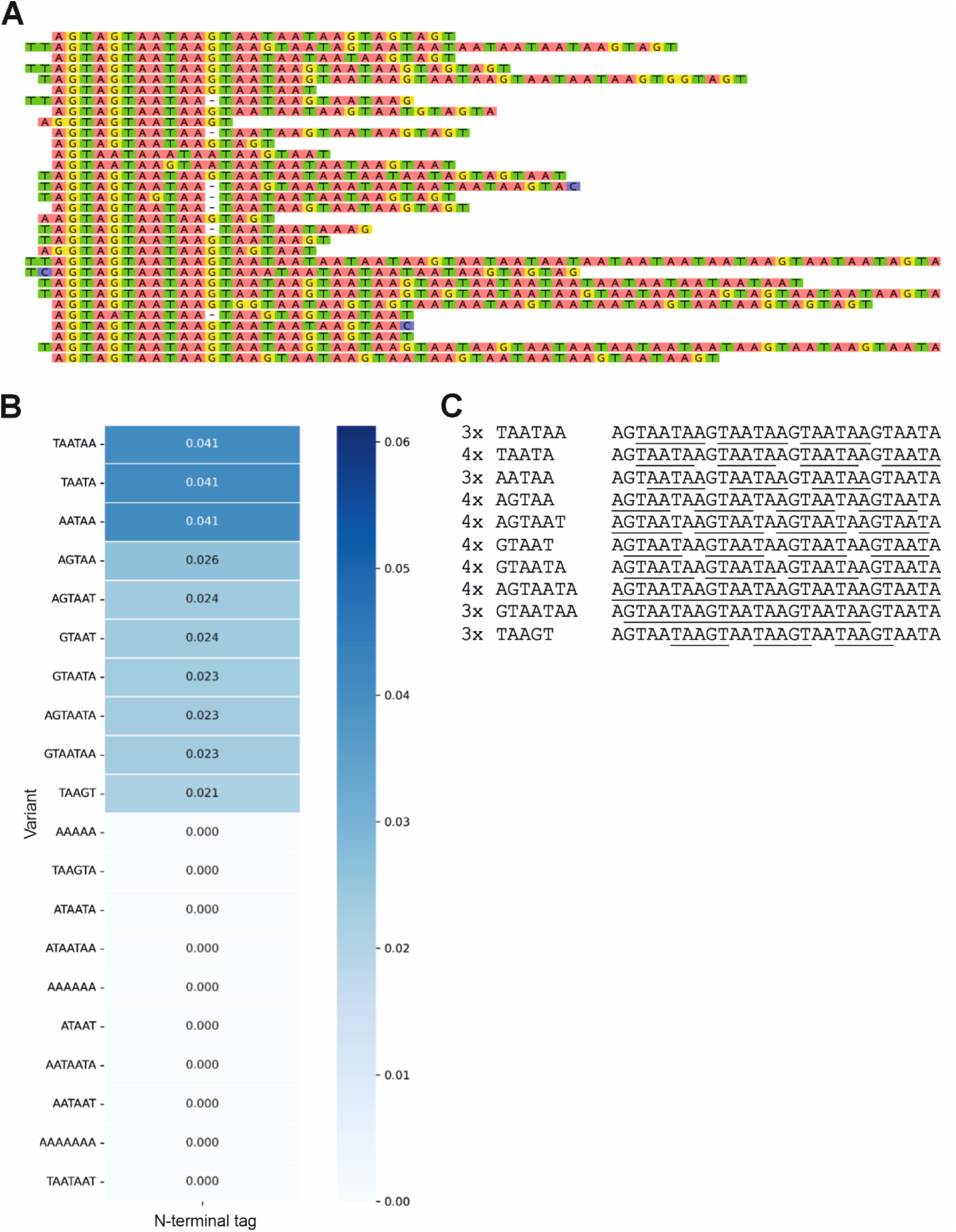
Identification of a recurrent AGTAATA repeat motif in DRT10-dependent cDNA products. (**A**) Representative unmapped ssDNA sequencing reads from *in vitro* DRT10 cDNA synthesis reactions showing tandem AGTAATA repeat tracts within unmapped reads. (**B**) Frequency of the 5–7-nt motifs enriched by k-mer analysis (k = 7) among unmapped reads. All enriched motifs are overlapping fragments derived from tandem AGTAATA repeat tracts. (**C**) Schematic representation showing that the ten most enriched motifs identified are all contained as overlapping substrings within tandem AGTAATA repeat tracts. Based on this observation, we designed a hybridization probe containing four tandem copies of the complementary sequence to AGTAATA (5’-TATTACTTATTACTTATTACTTATTACT-3’).

To confirm the identity of the repeat unit, we performed k-mer analysis of unmapped reads, which consistently identified AGTAATA-derived motifs as the most enriched sequences (**Figure 7B-C**), establishing AGTAATA as the consensus repeat unit of DRT10-generated cDNA products. Based on this observation, we designed an oligonucleotide probe complementary to tandem AGTAATA repeats to assess whether these cDNA products are also generated *in vivo*.

Using the AGTAATA-complementary probe, we analyzed total DNA from *E. coli* K-12 MG1655 ΔRM strains expressing DRT10. A hybridization signal was detected in wild-type strains both in the absence and presence of Bas52 infection, with signal intensity increasing upon infection, while no signal was observed in the RT-inactive mutant under any condition (**Figures 8A–B**), confirming strict RT-dependence. This phage-induced accumulation of the repetitive cDNA was also observed following co-immunoprecipitation of 6×His-tagged RT (**Figure 8C**). Together, these observations suggest that phage infection enhances cDNA accumulation.

**Figure 8.**
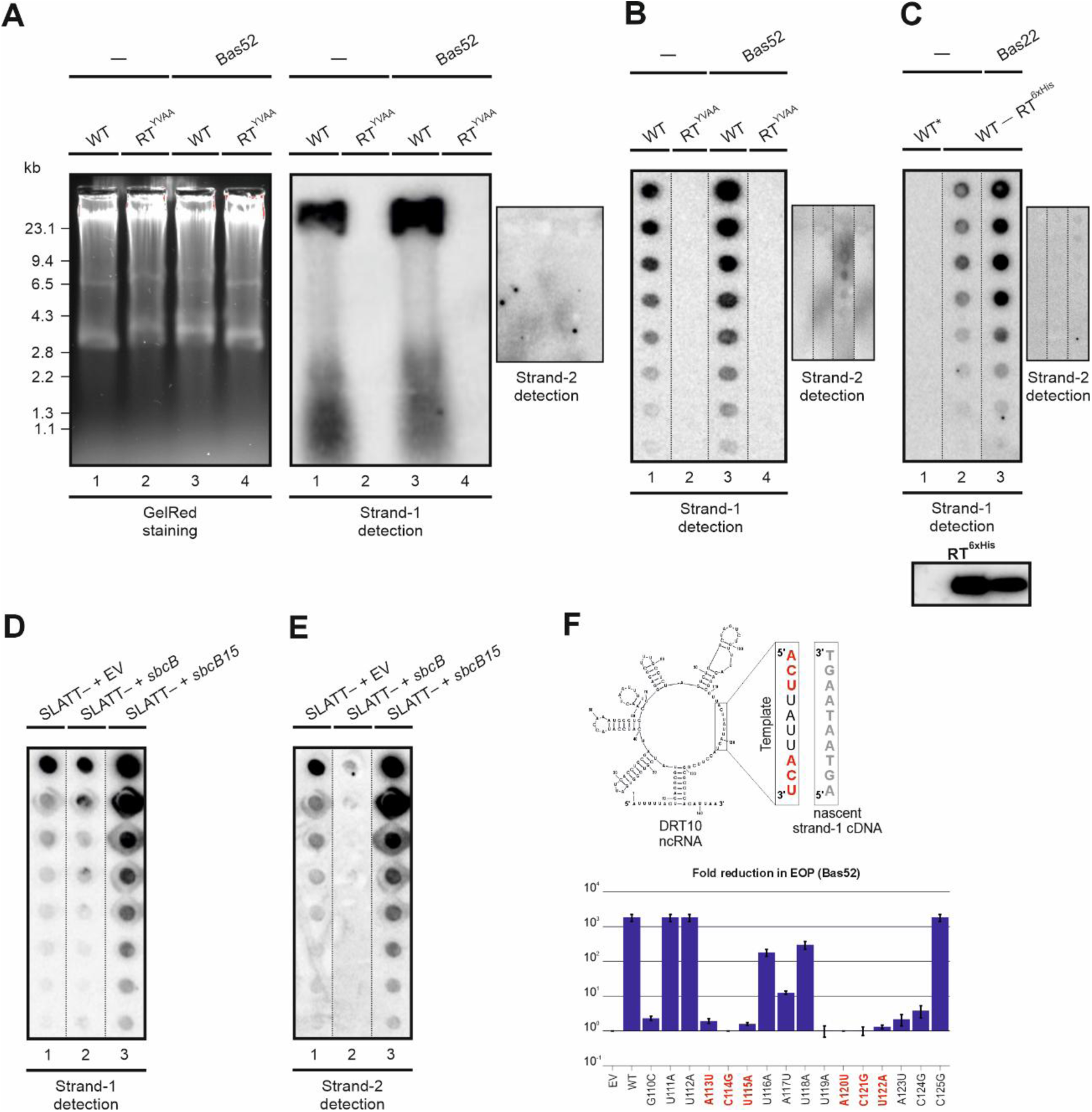
*In vivo* detection of DRT10-derived tandem-repeat cDNA and its ncRNA-encoded synthesis mechanism. (**A**) GelRed-stained agarose gel and Southern blot of total DNA from *E. coli* K-12 MG1655 ΔRM strains expressing either the wild-type DRT10 system (WT) or the RT-inactive mutant (RTYVAA), in the absence or presence of Bas52 phage infection, probed with a specific oligonucleotide targeting the first-strand cDNA. Adjacent small panel shows hybridization with a probe targeting the complementary second-strand cDNA. (**B**) Dot blot analysis of total DNA from the same strains and conditions as in panel **A,** probed with a specific oligonucleotide targeting the first-strand cDNA. Adjacent small panel shows hybridization with a probe targeting the complementary second-strand cDNA. (**C**) Dot blot analysis of DNA co-immunoprecipitated in *E. coli* K-12 MG1655 ΔRM strains expressing the control wild-type DRT10 system with the untagged RT (WT*) or the wild-type DRT10 system with the 6×His-tagged RT (WT) in the absence or presence of Bas22 phage infection, probed with a specific oligonucleotide targeting the first-strand cDNA. Bas22 phage was used in this assay because the 6×His-tagged RT construct confers only partial defense activity against Bas52, providing a wider dynamic range to detect cDNA accumulation-dependent differences. The Western blot confirming enrichment of the 6×His-tagged RT in the pull-down is shown below. Adjacent small panel shows hybridization with a probe targeting the complementary second-strand cDNA. (**D**) Dot blot analysis of total DNA from Δ*sbcB* strains expressing the SLATT− construct with the control pBB3 empty vector (EV), wild-type sbcB (pBB3_SbcB), or the sbcB15 variant (A183V) (pBB3_SbcB15), probed with a specific oligonucleotide targeting the first-strand cDNA. (**E**) Dot blot analysis of total DNA from the same strains and conditions as in panel D, probed with a specific oligonucleotide targeting the complementary second-strand cDNA. Equal amounts of total DNA were loaded per lane (OD600 = 0.5–0.6). The GelRed-stained gel is shown on the left, and the sizes of the GelRed-stained λII and φ29 DNA molecular weight markers are indicated on the left-hand side of panel A (**F**) Predicted secondary structure of the DRT10 ncRNA highlighting the 5’-ACUUAUUACU-3’ template region containing in red the two ACU triplets (positions 113–115 and 120–122), together with the nascent first-strand cDNA (5’-AGTAATAAGT-3’) shown in light gray. Mutated positions within the two ACU triplets are shown in red; and substitutions in the central region are shown in black. Fold reduction in Bas52 phage efficiency of plating (EOP) was determined by serial 10-fold dilution plaque assays comparing *E. coli* K-12 MG1655 ΔRM strains expressing the pUC57m empty vector (EV) with those expressing the wild-type DRT10 system (WT) or the indicated single-nucleotide substitution mutants. Data are representative of three independent replicates and are shown on the right-hand side of the pan

### SbcB selectively degrades DRT10-derived cDNA intermediates

To examine the role of SbcB in DRT10-derived cDNA accumulation, we analyzed cDNA levels in Δ*sbcB* strains expressing a SLATT-deficient DRT10 construct to maintain cell viability, complemented with either the empty pBB3 vector (EV), wild-type SbcB, or the catalytically impaired *sbcB15* allele. First-strand cDNA was detected in the absence of SbcB and at comparable levels upon co-expression of wild-type SbcB (**Figure 8D**), suggesting that this species is not efficiently degraded by SbcB. In contrast, co-expression of the dominant-negative *sbcB15* mutant led to markedly stronger cDNA accumulation (**Figure 8D**), consistent with a model in which this catalytically impaired variant shields the cDNA from degradation by other host exonucleases through non-productive substrate binding.

To further characterize the DRT10-derived cDNA products identified by hybridization, we performed strand-specific DNA sequencing on cells expressing a SLATT-deficient DRT10 construct in the *sbcB15* background, conditions that maximize cDNA accumulation, alongside the RT-inactive control (**Figure 9**).

**Figure 9.**
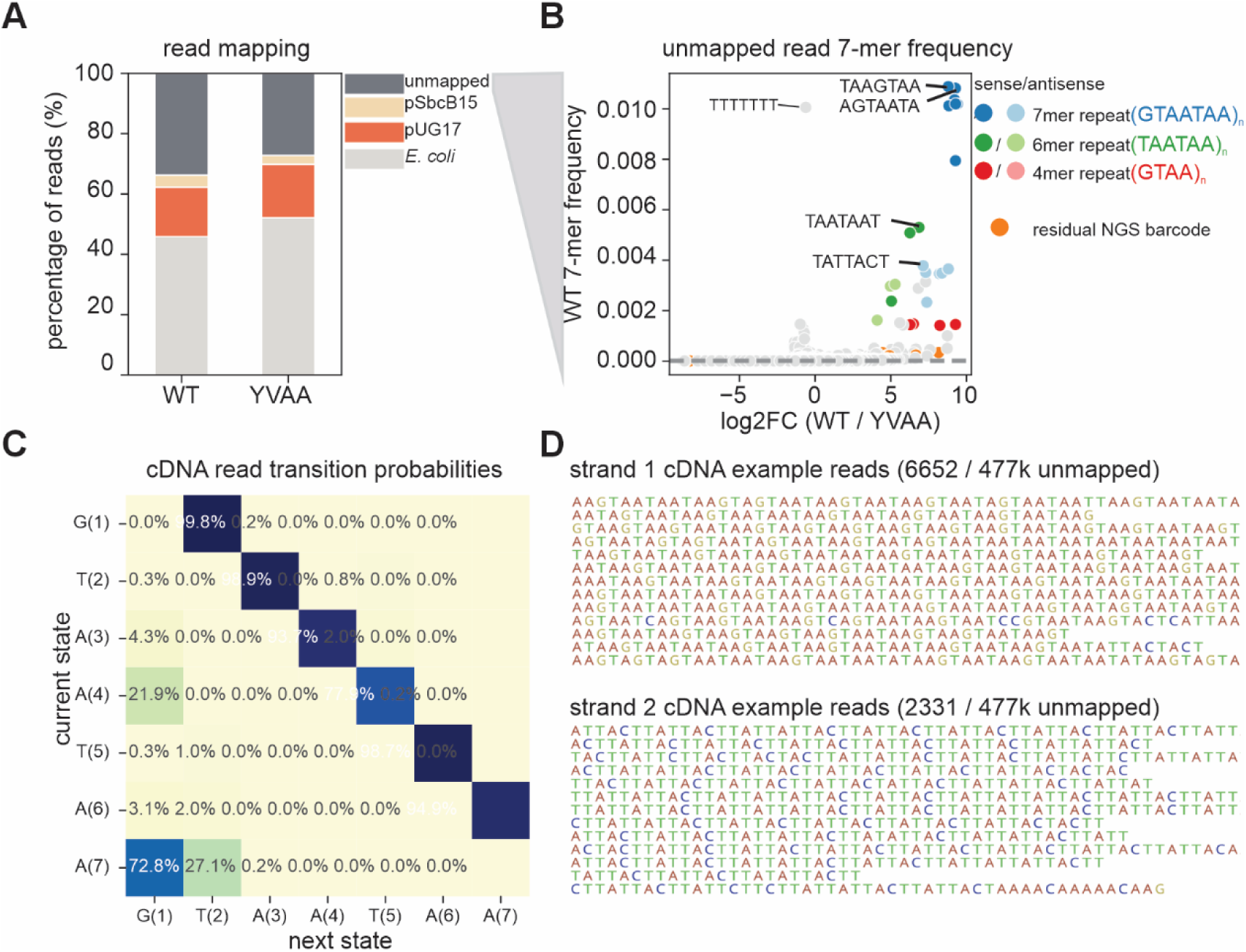
*In vivo* characterization of AGTAATA repeat-derived cDNA and strand distribution in DRT10-expressing cells. (**A**) Read-mapping representation for ssDNA-seq libraries generated from the SLATT-deficient construct (RT-active, WT) and the YVAA mutant (RT-inactive, YVAA), each expressed together with the *sbcB15* allele (A183V) in a Δ*sbcB* background. Bars are color-coded to denote mapping category (unmapped reads, dark gray; pBB3_sbcB15, salmon; DRT10-expression plasmid, orange; *E. coli* chromosome, light gray). (**B**) Unbiased 7-mer analysis of unmapped reads from RT-active (WT) and RT-inactive (YVAA) samples. Normalized 7-mer frequencies and enrichment values (RT-active/RT-inactive) are shown, with k-mers color-coded to distinguish cDNA-derived repeats (dark shades, sense orientation; light shades, antisense orientation) from residual untrimmed NGS adapter sequences (orange). Dark shades indicate sense orientation and light shades indicate antisense orientation. (**C**) Matrix transition-probability analysis of cDNA reads containing AGTAATAAGT in the RT-active sample. The repeat is represented in the GTAATAA rotated frame, which is equivalent to the AGTAATA repeat unit. (**D**) Representative ssDNA sequencing reads corresponding to strand 1 and the reverse-complement strand 2 detected in the RT-active sample. Numbers of reads assigned to each strand, as well as the total unmapped and total sequencing reads, are indicated below.

7-mer analysis of unmapped reads from the RT-active sample revealed strong enrichment of overlapping fragments of the AGTAATA tandem repeat, with no equivalent enrichment in the RT-inactive control (**Figures 9A–B**), confirming that the *in vivo* cDNA products consist of tandem AGTAATA repeats. Transition-probability analysis further revealed that while most repeat additions follow the expected AGTAATA pattern, a substantial fraction of reads exhibit imperfect junctions arising from occasional misalignment during template resetting (**Figure 9C**), indicating that tandem-repeat synthesis by DRT10 is processive but not perfectly precise.

Notably, the 7-mer TATTACT, corresponding to the complementary strand of the repeat, was also enriched in the RT-active sample (**Figure 9B**), and approximately one-third of repeat-derived cDNA reads mapped to the reverse-complement strand (**Figure 9D**). Although this fraction may partly reflect methodological biases inherent to cDIP-seq, these data are consistent with the synthesis of second-strand cDNA *in vivo*. To directly test this, we used a probe complementary to the first-strand AGTAATA repeat and detected a strong hybridization signal in *ΔsbcB* cells co-expressing *sbcB15*, whereas co-expression of wild-type SbcB abolished this signal (**Figure 8E**). Similarly to the first strand, the second-strand species was also stabilized by *sbcB15* (**Figure 8E**), demonstrating that both cDNA strands are protected by non-productive SbcB15 binding, while wild-type SbcB selectively degrades the second strand.

Together, these results indicate that, under conditions where SbcB is absent, both first-and second-strand cDNA species accumulate to similar levels. The preferential degradation of the second strand by SbcB suggests that it transiently exposes a free 3’ ssDNA end that is more accessible than the first strand 3’ end, which may remain partially protected in the RT active site and/or through duplex formation with the RNA template.

### Two ACU motifs within the ncRNA template program iterative tandem-repeat cDNA synthesis

To define how DRT10 programs synthesis of the AGTAATA repeat, we examined the ncRNA sequence and identified a single-stranded region of sequence 5’-ACU<u>UAUUACU</u>-3’, containing two ACU triplets that flank or are part of a sequence antisense to the AGTAATA repeat unit (underlined) (**Figure 8F**). Reverse transcription of this region would generate a nascent first strand terminating in AGT, whose complementarity to the downstream ACU triplet would allow iterative strand repositioning and repeat synthesis.

To test whether this architecture is required for repeat synthesis, we introduced single-nucleotide substitutions across the template loop, replacing each position with its complementary base. Substitutions within the central segment were generally well tolerated, whereas mutations disrupting either ACU triplet abolished phage defense (**Figure 8F**), demonstrating that both ACU motifs are essential for repeat formation and consistent with their proposed roles in template recognition and strand repositioning.

Together, these results support a model in which DRT10 generates AGTAATA tandem repeats through iterative cycles of reverse transcription, dissociation, and re-annealing of the nascent cDNA to the downstream ACU motif, which primes each subsequent round of synthesis (**Figure 8F**). Our sequencing data reveal that repeat addition is processive but not perfectly precise, with a substantial fraction of products exhibiting imperfect junctions arising from occasional misalignment during template resetting.

### A phage-encoded protein is required for DRT10-mediated defense activation

To identify phage determinants required for DRT10 activation, we isolated Bas52 escape mutants capable of evading DRT10-mediated defense. Two independent escape mutants were recovered, each carrying a single nonsynonymous substitution in the Bas52_0087 gene, resulting in D252V or Y351C amino acid changes. Both mutations enabled the phage to replicate in the presence of DRT10 (**Figure 10A**), identifying Bas52_0087 as a phage-encoded factor required for triggering DRT10-mediated defense.

**Figure 10.**
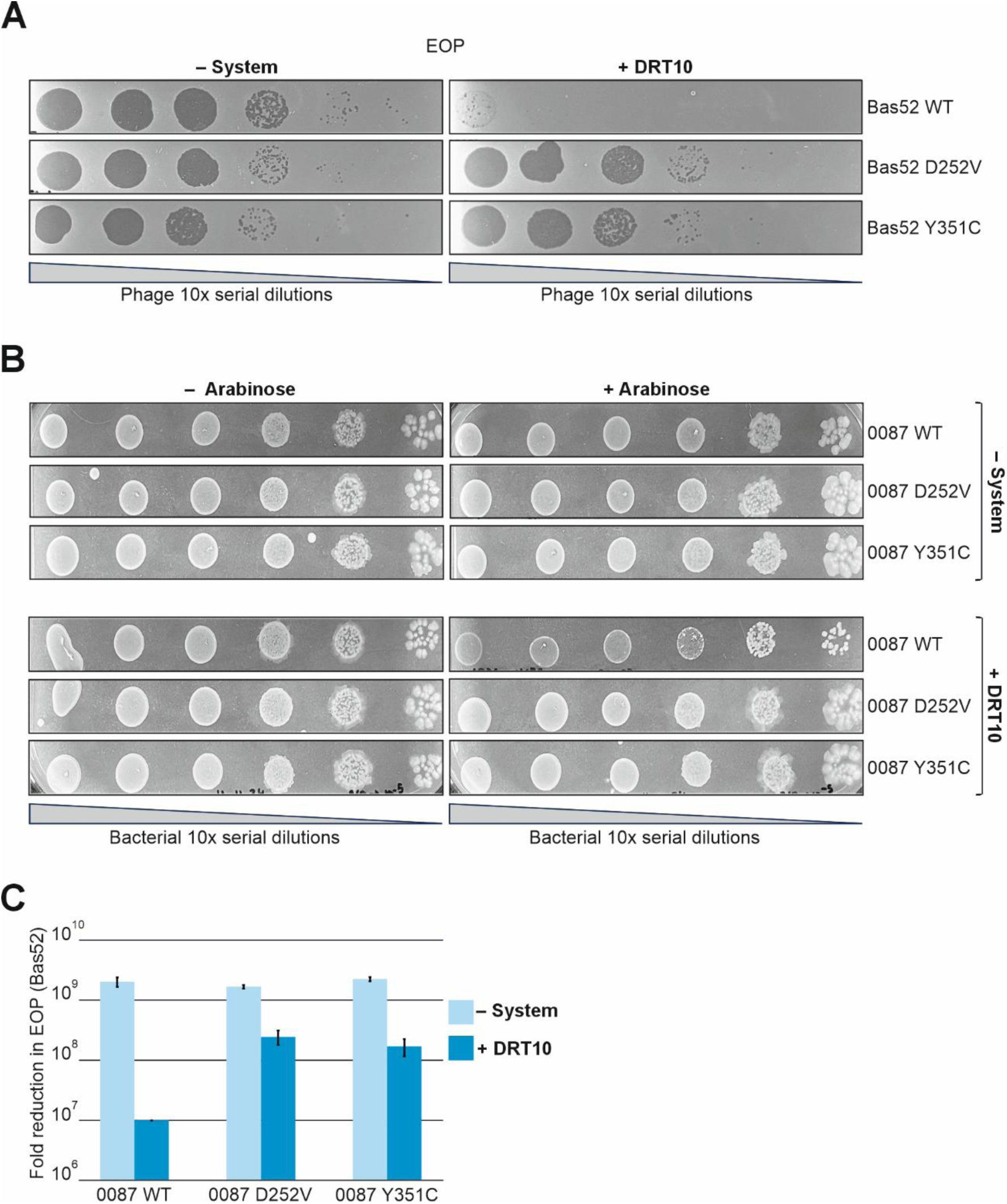
A phage-encoded protein is required for DRT10-mediated defense activation. (**A**) Serial 10-fold dilution plaque assays comparing the efficiency of plating (EOP) of the wild-type Bas52 phage (WT) or escape mutant phages (D_252_V and Y_351_C) on *E. coli* K-12 MG1655 ΔRM strains lacking (− system) or carrying (+ DRT10) the DRT10 system. (**B**) Cell viability of *E. coli* K-12 MG1655 ΔRM strains lacking or carrying the DRT10 system (expressed from a pUC-based plasmid) with either wild-type (WT) or mutant (D_252_V and Y_351_C) Bas52_0087 (ID: QXV84166.1) variants expressed from a pBAD33 plasmid, assessed by plating serial 10-fold dilutions without (− arabinose) and with (+ arabinose) induction of Bas52_0087 expression. (**C**) Fold reduction in EOP comparing Bas52 infectivity on *E. coli* K-12 MG1655 ΔRM strains lacking (− system) or carrying (+ DRT10) the DRT10 system with either wild-type (WT) or mutant (D_252_V and Y_351_C) Bas52_0087 variants expressed from a pBAD33 plasmid. Data are representative of three independent replicates.

To assess whether Bas52_0087 expression alone is sufficient to activate DRT10, we expressed the wild-type and mutant variants in *E. coli* in the presence or absence of the DRT10 system. Expression of Bas52_0087 did not affect host viability under either condition (**Figure 10B**), indicating that the phage protein requires additional factors present during infection to trigger DRT10-mediated defense.

To test whether Bas52_0087 modulates DRT10 activation during infection, we expressed the wild-type or escape mutant variants in the presence or absence of DRT10 and challenged the resulting strains with Bas52 phage. While wild-type Bas52_0087 expression markedly enhanced DRT10-mediated restriction, the D252V and Y351C variants showed substantially reduced ability to promote defense (**Figure 10C**), demonstrating that these substitutions impair the capacity of Bas52_0087 to activate DRT10 during phage infection.

Together, these results identify Bas52_0087 as a phage-encoded factor required for effective DRT10 activation, and suggest that additional phage-derived components present during infection contribute to full system engagement.

To investigate the interplay between SbcB, phage infection, and DRT10 activity, we overexpressed *sbcB* in cells carrying the DRT10 system and challenged them with Bas52 phage. Overexpression of SbcB attenuated DRT10-mediated restriction, as evidenced by an increased EOP relative to the empty vector control (**Figure S6A**). These findings demonstrate that ExoI dosage negatively modulates DRT10 defense activity during phage infection, establishing a direct link between host DNA processing capacity and the efficiency of DRT10-mediated immunity.

Given the connection between SbcB and DRT10 defense, we sought to further characterize the phage-encoded trigger Bas52_0087. Structural prediction using AlphaFold2 revealed an SSB-like architecture featuring two OB-fold domains and a C-terminal disordered region (**Figure S6B**). Notably, the C-terminal region harbors a conserved motif (FDDDIPF) closely resembling the C-terminal tip (C-tip) of *E. coli* SSB (**Figure S6B**), a sequence known to mediate interactions with multiple host DNA-processing factors including SbcB (Lu *et al*., 2011; Korada *et al*., 2013; Bianco and Lyubchenko, 2017; Pipalović *et al*., 2024). Furthermore, mapping the escape mutant substitutions D252V and Y351C onto the predicted structure placed both residues at the putative oligomerization interface of Bas52_0087 (**Figure S6C**), suggesting that oligomerization is required for its ability to trigger DRT10. Together, the SSB-like structure and the presence of a conserved C-terminal motif, raises the possibility that this phage protein may interface with host DNA-processing pathways.

## Discussion

Here we establish UG17 as a tripartite reverse transcriptase-dependent defense system, and provide a comprehensive characterization of its subtype diversity, genetic organization, and activity. We uncover a mechanistically distinct class-2 RT-based immune pathway built around a structured ncRNA, a SLATT effector, and a reverse transcriptase that synthesizes a discrete repeat-containing ssDNA species. The accumulation of this DNA intermediate is tightly controlled by the host exonuclease SbcB and a phage-encoded SSB-like trigger, revealing a defense strategy that integrates RT-driven DNA synthesis with host DNA-processing pathways.

We also systematically classified UG17 systems into three RT-defined subtypes with distinct gene architectures, taxonomic distributions, and subtype-specific ncRNAs, a diversity not fully captured in previous analyses. Phylogenies reconstructed from trimmed RT catalytic cores do not recover these subtypes as monophyletic, indicating that diversification primarily involves regions outside the conserved polymerase core, including gene architecture, ncRNA structure, and non-core RT domains. In contrast, SLATT proteins remain comparatively conserved, consistent with their role as a broadly maintained effector module. The restricted distribution of certain subtypes, such as the Bacillota-confined UG17-C lineage, further suggests host-associated specialization and raises the possibility that subtype-specific ncRNA features may reflect adaptation to distinct phage pressures.

The finding that the SL1-3 compensatory double mutant restores ncRNA accumulation yet fails to support RT-ncRNA association reveals an unexpected dissociation between RNA stability and functional complex formation. This result implies that the RT recognizes specific three-dimensional features of the ncRNA that are disrupted by the SL1-3 mutations despite restoration of base pairing, a level of structural specificity reminiscent of the stringent RNA recognition mechanisms described for other ncRNA-dependent RT systems such as DRT2 and DRT9. The identity of these higher-order structural determinants remains to be defined, but their existence suggests that the ncRNA functions not merely as a scaffold but as an active participant in RT recruitment and complex assembly. Once productively assembled, this RT-ncRNA complex drives a mechanistically distinct mode of DNA synthesis whose products are central to DRT10 immune function.

The discovery of tandem 7 nt repeat ssDNA generated by DRT10 adds new weight to the emerging trend that class 2 UG reverse transcriptases perform iterative repeat synthesis using single templates in ncRNAs. In all cases described so far, repeat synthesis is enabled by repeated motifs within the templates themselves. For DRT3 (Deng *et al*., 2026) and DRT9 (tang *et al*., 2025a), where their respective UG3 and UG28 reverse transcriptases synthesise relatively low-complexity poly(dTdG) or poly(dA) cDNA repeats, the templating ncRNA is inherently repetitive (e.g. ACACAC for DRT3; UUUU for DRT9), probably allowing slippage of the nascent cDNA. For DRT2 and DRT10, the 120 nt or 7 nt templates in the ncRNA contain trinucleotide repeats on each end of the template (“AUC” for DRT10, “ACA” for DRT2), allowing for slippage and precise repositioning once the template has finished one cycle of reverse transcription. In this way, the repeat synthesis mechanisms for DRT2 and DRT10 are reminiscent of telomerase, where two “CUAAC” pentamers flank the 6 nt template, leading to ‘repeat addition processivity’ (Wu *et al*., 2017; Nguyen, 2021). Based on this trend, we predict that further members of the class 2 UG RT family might also use ncRNA to template repeat synthesis with diverse possible template size.

We found that the DRT10 repetitive cDNA is constitutively synthesized under basal conditions and further accumulates upon phage infection, a behavior that contrasts with that reported for a related DRT10 homolog, in which cDNA abundance decreases upon infection (Tang *et al*., 2025b). This divergence may reflect differences between homologs rather than a fundamental mechanistic distinction, and underscores the importance of characterizing multiple representatives within a DRT family. We were also able to detect second-strand cDNA species when the exonuclease SbcB was inactivated.The differential accumulation of first- and second-strand cDNA species under different conditions reveals a previously unrecognized regulatory layer in DRT10 immunity. We suggest that SbcB selectively degrades the second strand, as demonstrated by its accumulation in the sbcB15 background and its absence when wild-type SbcB is present. Upon phage infection, first-strand accumulation is enhanced. Together, these observations suggest that the two cDNA species are differentially regulated, though the mechanistic basis of this asymmetry and its relationship to SLATT activation remain to be established.

The phage-encoded trigger Bas52_0087 provides a mechanistic link between phage infection and DRT10 activation. Structural predictions reveal an SSB-like architecture with a conserved C-terminal tip (C-tip) motif known to mediate interactions with host ssDNA-processing factors including SbcB, raising the intriguing possibility that Bas52_0087 may compete with or displace SbcB from ssDNA substrates during infection. The mapping of escape substitutions to a predicted oligomerization interface further suggests that functional multimerization of Bas52_0087 is required for its activating role. If Bas52_0087 indeed interferes with SbcB activity during infection, this could explain the enhanced first-strand accumulation observed upon phage challenge — not through inhibition of second-strand degradation per se, but by altering the overall ssDNA-processing landscape in ways that favor DRT10 engagement. These considerations suggest a model in which the phage trigger and host SbcB act as opposing regulators of DRT10 activation, whose relative activities determine the threshold for immune engagement.

Taken together, our findings support an emerging model in which DRT10 operates as a constitutive surveillance system whose activation threshold is jointly controlled by host DNA-processing capacity and phage-encoded factors, with SLATT potentially responding to different cDNA configurations depending on cellular context. How SLATT detects and responds to these signals, whether through direct recognition of the repeat-containing DNA or indirectly through downstream cellular perturbations, remains a central open question. Resolving the structural basis of the RT-ncRNA-cDNA complex, together with a mechanistic characterization of SLATT activation, will be essential to fully understand how DRT10 integrates constitutive DNA synthesis with context-dependent immune engagement.

## Acknowledgments

We thank Ascensión Martos Tejera for technical assistance.

## Author contributions

E.S.N. performed the experimental work and contributed to writing, reviewing, and editing the manuscript. V.M. contributed to experimental work and plasmid constructs.

C.W. performed the sequencing of cDNA products. D.C. contributed to initial characterization of the SbcB control of the DRT10 system. M.E.W. and F.M.A. supervised the study and the experimental work and contributed to manuscript writing.

N.T. supervised the study, conducted the ncRNA identification, structural and phylogenetic analyses, and contributed to writing, reviewing, and editing the manuscript.

## References

Baba, T., Ara, T., Hasegawa, M., Takai, Y., Okumura, Y., Baba, M., Datsenko, K. A., Tomita, M., Wanner, B. L., & Mori, H. (2006). Construction of *Escherichia coli* K-12 in-frame, single-gene knockout mutants: the Keio collection. Molecular Systems Biology, 2, 2006.0008. 10.1038/msb4100050

Baltimore, D. (1970). RNA-dependent DNA polymerase in virions of RNA tumour viruses. Nature, 226(5252), 1209–1211. 10.1038/2261209a0

Bianco, P. R., & Lyubchenko, Y. L. (2017). SSB and the RecG DNA helicase: an intimate association to rescue a stalled replication fork. Protein Science, 26(3), 638–649. 10.1002/pro.3114

Bouras, G., Grigson, S. L., Papudeshi, B., Mallawaarachchi, V., & Edwards, R. A. (2024). Prokaryotic virus remote homology detection and functional annotation using language models. PLOS Computational Biology, 20(11), e1012645. 10.1371/journal.pcbi.1012645

Burroughs, A. M., Zhang, D., Schäffer, D. E., Iyer, L. M., & Aravind, L. (2015). Comparative genomic analyses reveal a vast, novel network of nucleotide-centric systems in biological conflicts, immunity and signaling. Nucleic Acids Research, 43(22), 10633–10654. 10.1093/nar/gkv1267

Cabanes, D., Boistard, P., & Batut, J. (2000). Symbiotic induction of pyruvate dehydrogenase genes from *Sinorhizobium meliloti*. Molecular Plant-Microbe Interactions (MPMI), 13(5), 483–493. 10.1094/MPMI.2000.13.5.483

Capella-Gutiérrez, S., Silla-Martínez, J. M., & Gabaldón, T. (2009). trimAl: a tool for automated alignment trimming in large-scale phylogenetic analyses. Bioinformatics, 25(15), 1972–1973. 10.1093/bioinformatics/btp348

del Val, C., Rivas, E., Torres-Quesada, O., Toro, N., & Jiménez-Zurdo, J. I. (2007). Identification of differentially expressed small non-coding RNAs in the legume endosymbiont *Sinorhizobium meliloti* by comparative genomics. Molecular Microbiology, 66(5), 1080–1091. 10.1111/j.1365-2958.2007.05978.x

Deng, P., Lee, H., Armijo, C., Wang, H., & Gao, A. (2026). Protein-templated synthesis of dinucleotide repeat DNA by an antiphage reverse transcriptase. *Science*, eaed1656. 10.1126/science.aed1656

Eddy, S. R. (2011). Accelerated profile HMM searches. PLOS Computational Biology, 7(10), e1002195. 10.1371/journal.pcbi.1002195

Fekete, R. A., Miller, M. J., & Chattoraj, D. K. (2003). Fluorescently labeled oligonucleotide extension: a rapid and quantitative protocol for primer extension. BioTechniques, 35(1), 90–98. 10.2144/03351rr01

Figiel, M., Gapińska, M., Czarnocki-Cieciura, M., Zajko, W., Sroka, M., Skowronek, K., & Nowotny, M. (2022). Mechanism of protein-primed template-independent DNA synthesis by Abi polymerases. Nucleic Acids Research, 50(17), 10026–10040. 10.1093/nar/gkac772

Fillol-Salom, A., Rostøl, J. T., Ojiogu, A. D., Chen, J., Douce, G., Humphrey, S., & Penadés, J. R. (2022). Bacteriophages benefit from mobilizing pathogenicity islands encoding immune systems against competitors. Cell, 185(17), 3248–3262.e20. 10.1016/j.cell.2022.07.014

Fu, L., Niu, B., Zhu, Z., Wu, S., & Li, W. (2012). CD-HIT: accelerated for clustering the next-generation sequencing data. Bioinformatics, 28(23), 3150–3152. 10.1093/bioinformatics/bts565

Gao, L., Altae-Tran, H., Böhning, F., Makarova, K. S., Segel, M., Schmid-Burgk, J. L., Koob, J., Wolf, Y. I., Koonin, E. V., & Zhang, F. (2020). Diverse enzymatic activities mediate antiviral immunity in prokaryotes. Science, 369(6507), 1077–1084. 10.1126/science.aba0372

Gapińska, M., Zajko, W., Skowronek, K., Figiel, M., Krawczyk, P. S., Egorov, A. A., Dziembowski, A., Johansson, M. J. O., & Nowotny, M. (2024). Structure-functional characterization of Lactococcus AbiA phage defense system. Nucleic Acids Research, 52(8), 4723–4738. 10.1093/nar/gkae230

García-Tomsig, N. I., García-Rodríguez, F. M., Guedes-García, S. K., Millán, V., Becker, A., Robledo, M., & Jiménez-Zurdo, J. I. (2023). A double-negative feedback loop between NtrBC and a small RNA rewires nitrogen metabolism in legume symbionts. mBio, 14(6), e0200323. 10.1128/mbio.02003-23

González-Delgado, A., Mestre, M. R., Martínez-Abarca, F., & Toro, N. (2021). Prokaryotic reverse transcriptases: from retroelements to specialized defense systems. FEMS Microbiology Reviews, 45(6), fuab025. 10.1093/femsre/fuab025

Han, J., Liu, B., Tang, J., Zhang, S., Wang, X., Li, X., Zhang, Q., Liu, Z., Wang, W., Liu, Y., Zhou, R., Yin, H., Wei, Y., Li, Z., Zhang, M., Deng, Z., & Zhang, H. (2025). Non-coding RNA mediates the defense-associated reverse transcriptase (DRT) anti-phage oligomerization transition. The EMBO Journal, 44(19), 5429–5442. 10.1038/s44318-025-00544-8

Jumper, J., Evans, R., Pritzel, A., Green, T., Figurnov, M., Ronneberger, O., et al. (2021). Highly accurate protein structure prediction with AlphaFold. Nature, 596, 583–589. 10.1038/s41586-021-03819-2

Katoh, K., & Standley, D. M. (2013). MAFFT multiple sequence alignment software version 7: improvements in performance and usability. Molecular Biology and Evolution, 30(4), 772–780. 10.1093/molbev/mst010

Khan, A. G., Rojas-Montero, M., González-Delgado, A., Lopez, S. C., Fang, R. F., Crawford, K. D., et al. (2025). An experimental census of retrons for DNA production and genome editing. Nature Biotechnology, 43, 914–922. 10.1038/s41587-024-02384-z

Korada, S. K., Johns, T. D., Smith, C. E., Jones, N. D., McCabe, K. A., & Bell, C. E. (2013). Crystal structures of *Escherichia coli* exonuclease I in complex with single-stranded DNA provide insights into the mechanism of processive digestion. Nucleic Acids Research, 41(11), 5887–5897. 10.1093/nar/gkt278

Kropinski, A. M., Mazzocco, A., Waddell, T. E., Lingohr, E., & Johnson, R. P. (2009). Enumeration of bacteriophages by double agar overlay plaque assay. In Methods in Molecular Biology (Vol. 501, pp. 69–76). Humana Press. 10.1007/978-1-60327-164-6_7

Langmead, B., & Salzberg, S. L. (2012). Fast gapped-read alignment with Bowtie 2. Nature Methods, 9(4), 357–359. 10.1038/nmeth.1923

Lu, D., Myers, A. R., George, N. P., & Keck, J. L. (2011). Mechanism of Exonuclease I stimulation by the single-stranded DNA-binding protein. Nucleic Acids Research, 39(15), 6536–6545. 10.1093/nar/gkr315

Maffei, E., Shaidullina, A., Burkolter, M., Heyer, Y., Estermann, F., Druelle, V., Sauer, P., Willi, L., Michaelis, S., Hilbi, H., Thaler, D. S., & Harms, A. (2021). Systematic exploration of *Escherichia coli* phage-host interactions with the BASEL phage collection. PLOS Biology, 19(11), e3001424. 10.1371/journal.pbio.3001424

Martin, M. (2011). Cutadapt removes adapter sequences from high-throughput sequencing reads. EMBnet.journal, 17(1), 10–12. 10.14806/ej.17.1.200

Mestre, M. R., Gao, L. A., Shah, S. A., López-Beltrán, A., González-Delgado, A., Martínez-Abarca, F., Iranzo, J., Redrejo-Rodríguez, M., Zhang, F., & Toro, N. (2022). UG/Abi: a highly diverse family of prokaryotic reverse transcriptases associated with defense functions. Nucleic Acids Research, 50(11), 6084–6101. 10.1093/nar/gkac467

Minh, B. Q., Schmidt, H. A., Chernomor, O., Schrempf, D., Woodhams, M. D., von Haeseler, A., & Lanfear, R. (2020). IQ-TREE 2: New models and efficient methods for phylogenetic inference in the genomic era. Molecular Biology and Evolution, 37(5), 1530–1534. 10.1093/molbev/msaa015

Millman, A., Bernheim, A., Stokar-Avihail, A., Fedorenko, T., Voichek, M., Leavitt, A., Oppenheimer-Shaanan, Y., & Sorek, R. (2020). Bacterial retrons function in anti-phage defense. Cell, 183(6), 1551–1561.e12. 10.1016/j.cell.2020.09.065

Molina-Sánchez, M. D., Martínez-Abarca, F., & Toro, N. (2010). Structural features in the C-terminal region of the *Sinorhizobium meliloti* RmInt1 group II intron-encoded protein contribute to its maturase and intron DNA-insertion function. The FEBS Journal, 277(1), 244–254. 10.1111/j.1742-4658.2009.07478.x

Muñoz-Adelantado, E., San Filippo, J., Martínez-Abarca, F., García-Rodríguez, F. M., Lambowitz, A. M., & Toro, N. (2003). Mobility of the *Sinorhizobium meliloti* group II intron RmInt1 occurs by reverse splicing into DNA, but requires an unknown reverse transcriptase priming mechanism. Journal of Molecular Biology, 327(5), 931–943. 10.1016/s0022-2836(03)00208-0

Nguyen, T. H. D. (2021). Structural biology of human telomerase: progress and prospects. Biochemical Society Transactions, 49(5), 1927–1939. 10.1042/BST20200042

Pipalović, G., Filić, Ž., Ćehić, M., Paradžik, T., Zahradka, K., Crnolatac, I., & Vujaklija, D. (2024). Impact of C-terminal domains of paralogous single-stranded DNA binding proteins from *Streptomyces coelicolor* on their biophysical properties and biological functions. International Journal of Biological Macromolecules, 268(Pt 1), 131544. 10.1016/j.ijbiomac.2024.131544

Price, M. N., Dehal, P. S., & Arkin, A. P. (2010). FastTree 2--approximately maximum-likelihood trees for large alignments. PLOS ONE, 5(3), e9490. 10.1371/journal.pone.0009490

Reuter, J. S., & Mathews, D. H. (2010). RNAstructure: software for RNA secondary structure prediction and analysis. BMC Bioinformatics, 11, 129. 10.1186/1471-2105-11-129

Rivas, E. (2020). RNA structure prediction using positive and negative evolutionary information. PLOS Computational Biology, 16(10), e1008387. 10.1371/journal.pcbi.1008387

Rivas, E., Clements, J., & Eddy, S. R. (2020). Estimating the power of sequence covariation for detecting conserved RNA structure. Bioinformatics, 36(10), 3072–3076. 10.1093/bioinformatics/btaa080

Rousset, F., Cabezas-Caballero, J., Piastra-Facon, F., Fernández-Rodríguez, J., Clermont, O., Denamur, E., Rocha, E. P. C., & Bikard, D. (2021). The impact of genetic diversity on gene essentiality within the *Escherichia coli* species. Nature Microbiology, 6(3), 301–312. 10.1038/s41564-020-00839-y

Schuster, C. F., & Bertram, R. (2014). Fluorescence based primer extension technique to determine transcriptional starting points and cleavage sites of RNases in vivo. Journal of Visualized Experiments*, (*92), e52134. 10.3791/52134

Simon, D. M., & Zimmerly, S. (2008). A diversity of uncharacterized reverse transcriptases in bacteria. Nucleic Acids Research, 36(22), 7219–7229. 10.1093/nar/gkn867

Song, X. Y., Xia, Y., Zhang, J. T., Liu, Y. J., Qi, H., Wei, X. Y., Hu, H., Xia, Y., Liu, X., Ma, Y. F., & Jia, N. (2025). Bacterial reverse transcriptase synthesizes long poly(A)-rich cDNA for antiphage defense. Science, 388(6753), eads4639. 10.1126/science.ads4639

Tang, S., Conte, V., Zhang, D. J., Žedaveinytė, R., Lampe, G. D., Wiegand, T., Tang, L. C., Wang, M., Walker, M. W. G., George, J. T., Berchowitz, L. E., Jovanovic, M., & Sternberg, S. H. (2024). De novo gene synthesis by an antiviral reverse transcriptase. Science, 386(6717), eadq0876. 10.1126/science.adq0876

Tang, S., Žedaveinytė, R., Burman, N., Pandey, S., Ramirez, J. L., Kulber, L. M., Wiegand, T., Wilkinson, R. A., Ma, Y., Zhang, D. J., Lampe, G. D., Berisa, M., Jovanovic, M., Wiedenheft, B., & Sternberg, S. H. (2025a). Protein-primed homopolymer synthesis by an antiviral reverse transcriptase. Nature, 643(8074), 1352–1362. 10.1038/s41586-025-09179-5

Tang, S., Ramirez, J. L., Rodríguez Mestre, M., Zhang, D. J., Wang, M., Wiegand, T., Ma, Y., Jovanovic, M., Pinilla-Redondo, R., & Sternberg, S. H. (2025b). Antiviral reverse transcriptases reveal the evolutionary origin of telomerase. bioRxiv preprint. 10.1101/2025.10.16.682844

Temin, H. M., & Mizutani, S. (1970). RNA-dependent DNA polymerase in virions of Rous sarcoma virus. Nature, 226(5252), 1211–1213. 10.1038/2261211a0

Thoms, B., Borchers, I., & Wackernagel, W. (2008). Effects of single-strand DNases ExoI, RecJ, ExoVII, and SbcCD on homologous recombination of recBCD+ strains of *Escherichia coli* and roles of SbcB15 and XonA2 ExoI mutant enzymes. Journal of Bacteriology, 190(1), 179–192. 10.1128/JB.01052-07

Toro, N., Martínez-Abarca, F., Mestre, M. R., & González-Delgado, A. (2019). Multiple origins of reverse transcriptases linked to CRISPR-Cas systems. RNA Biology, 16(10), 1486–1493. 10.1080/15476286.2019.1639310

Toro, N., & Nisa-Martínez, R. (2014). Comprehensive phylogenetic analysis of bacterial reverse transcriptases. PLOS ONE, 9(11), e114083. 10.1371/journal.pone.0114083

Wang, Y., Wu, H., Li, J., Tian, Z., & Deng, Z. (2025). Cryo-EM structure of the bacterial anti-phage defense system DRT6. Biochemical and Biophysical Research Communications, 791, 152955. 10.1016/j.bbrc.2025.152955

Wilkinson, M. E., Li, D., Gao, A., Macrae, R. K., & Zhang, F. (2024). Phage-triggered reverse transcription assembles a toxic repetitive gene from a noncoding RNA. Science, 386(6717), eadq3977. 10.1126/science.adq3977

Will, S., Reiche, K., Hofacker, I.L., Stadler, P.F. & Backofen R (2007). Inferring Noncoding RNA Families and Classes by Means of Genome-Scale Structure-Based Clustering. PLoS Computational Biology, 3(*4*): e65. 10.1371/journal.pcbi.0030065

Wu, R. A., Upton, H. E., Vogan, J. M., & Collins, K. (2017). Telomerase mechanism of telomere synthesis. Annual Review of Biochemistry, 86, 439–460. 10.1146/annurev-biochem-061516-045019

Yee, T., Furuichi, T., Inouye, S., & Inouye, M. (1984). Multicopy single-stranded DNA isolated from a gram-negative bacterium, *Myxococcus xanthus*. Cell, 38(1), 203–209. 10.1016/0092-8674(84)90541-5

Yu, G., Smith, D. K., Zhu, H., Guan, Y., & Lam, T. T. Y. (2017). ggtree: an R package for visualization and annotation of phylogenetic trees with their covariates and related data. Methods in Ecology and Evolution, 8(1), 28–36. 10.1111/2041-210X.12628

Zimmerly, S., & Wu, L. (2015). An unexplored diversity of reverse transcriptases in bacteria. Microbiology Spectrum, 3(2), MDNA3-2014. 10.1128/microbiolspec.MDNA3-0058-2014

