## Supplementary Figures for "Host exonuclease SbcB and a phage-encoded SSB-like protein control activation of the DRT10 reverse transcriptase defense system"

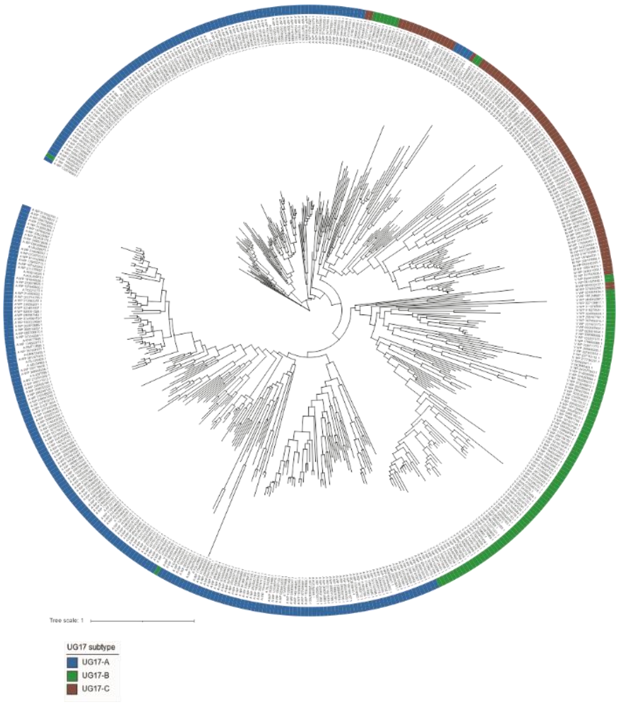


**Figure S1. Maximum-likelihood phylogenetic tree of RT proteins from the expanded UG17 dataset inferred using IQ-TREE2.** Protein sequences (n = 484) were aligned with MAFFT and trimmed using trimAl (Capella-Gutiérrez et al., 2009) with a custom Python script to retain the conserved RT core prior to tree inference. The best-fitting amino acid substitution model was selected using ModelFinder, and branch support was assessed using ultrafast bootstrap and SH-aLRT tests (1,000 replicates each). The tree is shown unrooted. Sequences are coloured according to their UG17 subtype assignment (UG17-A, UG17-B, and UG17-C). When analysis is restricted to the conserved RT core, the three subtypes are not recovered as strictly monophyletic clades.

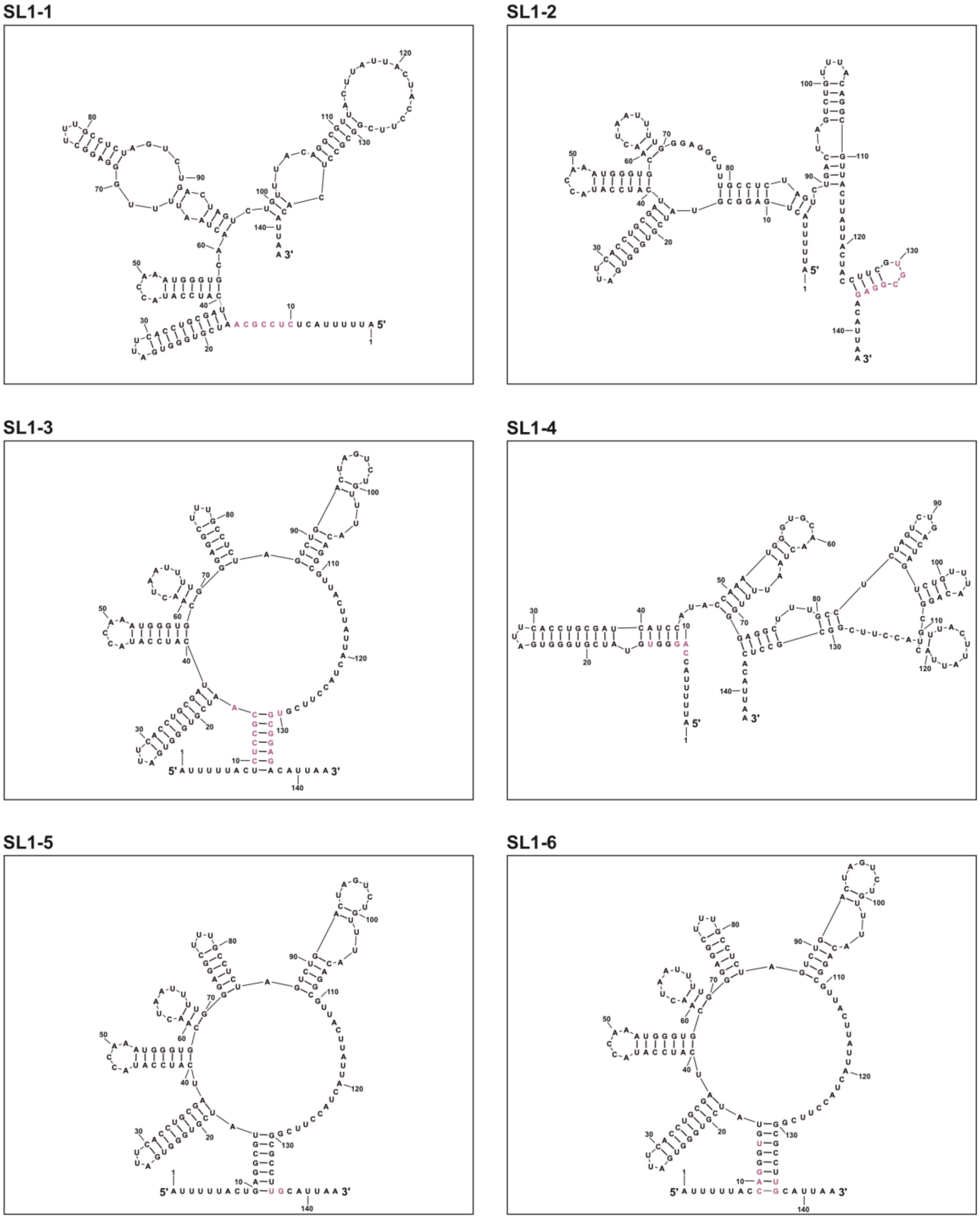


**Figure S2. Predicted structures of UG17-A mutant ncRNAs.** Predicted secondary structures of the ncRNA mutants in the basal SL1 stem (SL1-1 to SL1-6) used in the different assays. All predictions were obtained with RNAstructure (Reuter & Mathews, 2010).

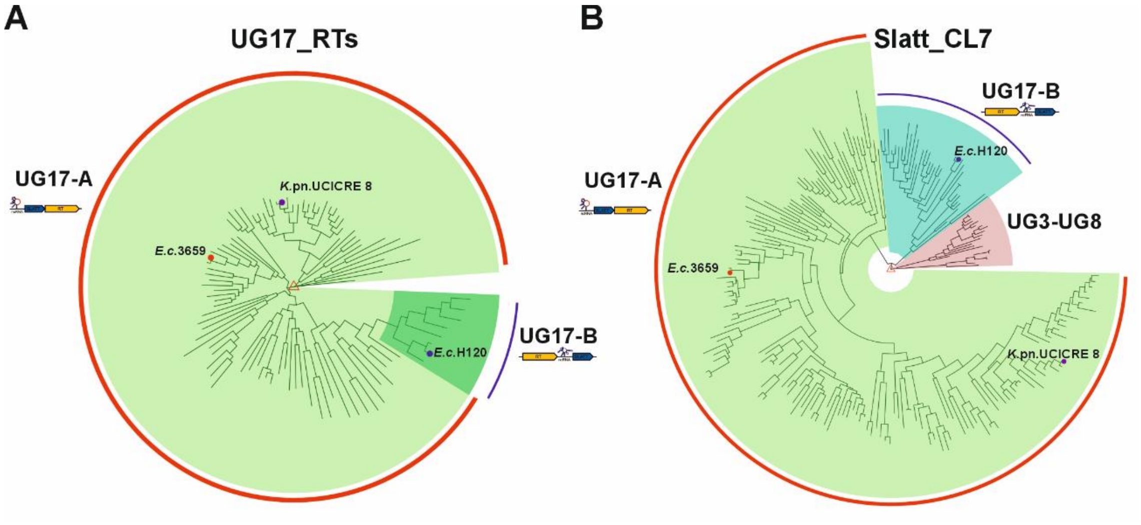


**Figure S3. UG17 systems previously misclassified as retrons.** (**A**) Phylogenetic tree of UG17-associated reverse transcriptases (RTs). (**B**) Phylogenetic tree of UG17-associated SLATT proteins. For UG17 systems, only SLATT proteins from cluster 7 (CL7; Mestre *et al*., 2022) were included. The trees feature RTs and SLATT proteins of *E. coli* H120 (Rousset *et al*., 2021**)**, marked with a blue dot, and the PICI-immune system from *Klebsiella pneumoniae* (KpCIUCICRE8, Fillol-Salom *et al*., 2022), marked with a purple dot, both previously misclassified as retrons. RT and SLATT proteins from the UG17-A E3659 system are also indicated with a red dot. SLATT proteins from cluster 7 of the UG3-UG8 (DRT3) system were included as an outgroup. Trees were generated using FastTree based on a MAFFT alignment, and the final figure was prepared using the *ggtree* package (Yu *et al*., 2017) in R.

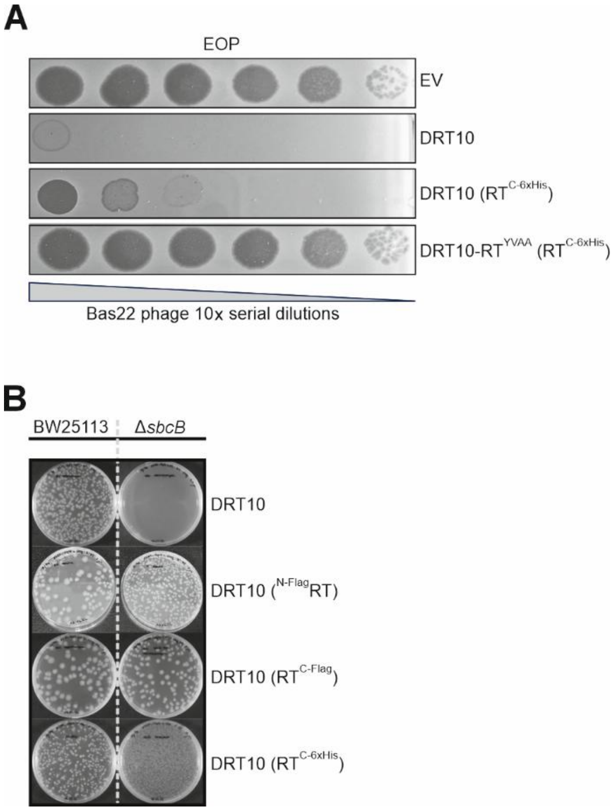


**Figure S4. Phenotypes of DRT10 RT constructs.** (**A**) Serial 10-fold dilution plaque assays comparing the efficiency of plating (EOP) of phage Bas22 on *E. coli* K-12 MG1655 ΔRM strains expressing the control pUC57m empty vector (EV), the wild-type DRT10 system with the untagged RT, the wild-type DRT10 system with the C-terminally 6×His-tagged RT ($RT^{C\text{-}6\times His}$), or the RT mutant (YVAA) carrying a C-terminal 6×His-tagged RT ($RT^{C\text{-}6\times His}$). Bas22 was selected for this assay because the 6×His-tagged RT construct confers only partial defense activity against Bas52, whereas it retains sufficient activity against Bas22, providing a wider dynamic range to detect phenotypic differences. (**B**) Transformation assays in *E. coli* K-12 BW25113 and its isogenic Δ*sbcB* derivative with the wild-type DRT10 system with the untagged RT, the DRT10 system with the Flag-tagged RT at either terminus ($^{N\text{-}Flag}RT$ and $RT^{C\text{-}Flag}$), or the DRT10 system with the C-terminally 6×His-tagged RT ($RT^{C\text{-}6\times His}$). Flag tagging at either terminus abolished DRT10-associated toxicity in the ΔsbcB background, whereas the C-terminal 6×His tag retained partial activity, supporting its use in subsequent co-immunoprecipitation experiments.

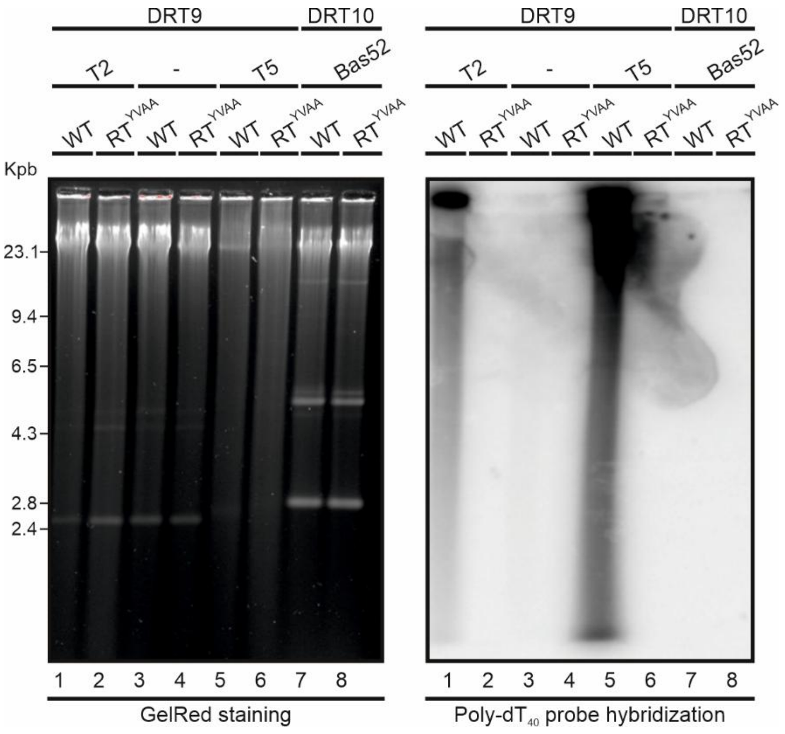


**Figure S5**. **Detection of poly-dA products.** GelRed-stained agarose gel and Southern blot of total DNA from *E. coli* K-12 MG1655 strains expressing either the wild-type *Sen*DRT9 system (WT) or the RT-inactive mutant (RT$^{YVAA}$), in the absence or presence of phage T2 or T5 infection, and from *E. coli* K-12 MG1655 ΔRM strains expressing either the wild-type DRT10 system (WT) or the RT-inactive mutant (RT$^{YVAA}$) upon Bas52 phage infection. Equal amounts of total DNA were loaded per lane ($OD_{600}$ = 0.5–0.6). The blot was probed with a specific oligonucleotide targeting the poly-dA products (5’-TTTTTTTTTTTTTTTTTTTTTTTTTTTTTTTTTTTTTTTT-3’). The GelRed-stained gel is shown on the left, and the sizes of the GelRed-stained λII and φ29 DNA molecular weight markers are indicated on the left-hand side of the gel.

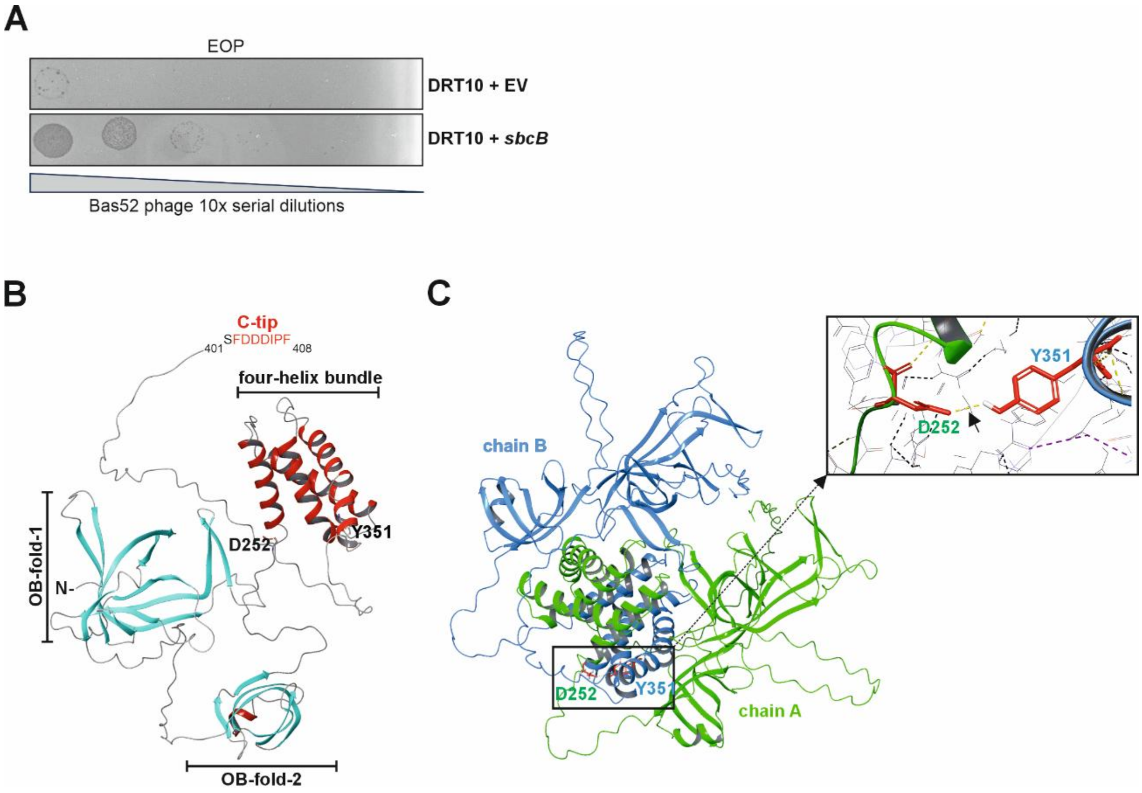


**Figure S6. SbcB modulates DRT10 defense and the phage trigger encodes a predicted SSB-like protein.** (**A**) Serial 10-fold dilution plaque assays comparing the efficiency of plating (EOP) of Bas52 phage on *E. coli* K-12 MG1655 ΔRM strains expressing the wild-type DRT10 system with either the empty pBB3 vector (EV) or the pBB3_SbcB plasmid containing the *sbcB* gene. Data are representative of three independent replicates. (**B**) Predicted three-dimensional structure of Bas52_0087, functionally annotated as an SSB-like protein by Phold. The structure comprises two OB-fold domains (OB-fold-1 and OB-fold-2, blue), a four-helix bundle and C-terminal tip (C-tip, red), and a C-terminal disordered region (grey). The residues mutated in Bas52 escape mutants (D252V, green; Y351C, blue) are shown. (**C**) Predicted spatial arrangement of residues $D_{252}V$ (green) and $Y_{351}C$ (blue) at the oligomerization interface of the Bas52_0087 protein. All structure predictions were obtained using AlphaFold2 (Jumper et al., 2021).
