## Supplementary material for "Host exonuclease SbcB and a phage-encoded SSB-like protein control activation of the DRT10 reverse transcriptase defense system": Table S1

| DNA Oligonucleotide | 5’-sequence-3’ | Use |
|---|---|---|
| Flag_1_Fw | CAGATGTTAAGGATGCTGGTTACCTAGAGGTTCTCTTTC | Amplification of Flag tag fragment (used for RT domain tagging) |
| Flag_2_Rv | GTTGCTAATGAGCATAATTATTACTTGTCATCGTCATCC | |
| Flag_3_Rv | GTAACCAGCATCCTTAACATC | Amplification of UG17 locus fragment (inverse PCR for Flag-tagging) |
| Flag_4_Fw | TAATAATTATGCTCATTAGCAAC | |
| His_Fw-*Xho*I | GGGCTCGAGCACCACCACCACCACCACTAATAATTATGCTC ATTAGCAACA | Amplification of 6×His tag fragment (inverse PCR for RT domain tagging) |
| His_Rv-*Xho*I | GGGCTCGAGGTAACCAGCATCCTTAACATC | |
| 422_Fw-*EcoR*I | GGGGAATTCGGTGCGGTGTAGTGCGATT | Amplification of UG17 system fragment |
| Bas52_Fw-*Kpn*I | GGGGTACCAAGGAGACAAATAATGAGCGT | Amplification of *Bas52_0087* phage gene |
| Bas52_Rv-*Sal*I | CCCGTCGACTTATTAAAAAGGGATGTCATCATCA | |
| *sbcB*_Fw-*Spe*I | GGACTAGTCAGCAAACCCTCAGGAGTT | Amplification of *sbcB* and *sbcB15* genes ($A_{183}V$) |
| *sbcB*_Rv-*Kpn*I | GGGGGTACCTACCAGCGGCGGAGGCTT | Amplification of *sbcB* gene |
| *sbcB15*_Rv-*Xcm*I | GCTTTGCCATCGCAATAGTGGCGTACACATCAGCCATCACAT CGT | Amplification of s*bcB15* mutant gene |
| 13_Fw-*Kpn*I | GGGGGTACCCCGAAAGGCATTTACACAGA | Amplification of wild-type ncRNA |
| 622_Rv-*Sac*I | GGGGAGCTCAATCAAGATTAAGAAGGCGG | |
| Probe_anti-cDNA | TATTACTTATTACTTATTACTTATTACT | Detection of UG17 cDNA (blot hybridization) |
| Probe_poly-$dT_{40}$ | TTTTTTTTTTTTTTTTTTTTTTTTTTTTTTTTTTTTTTTT | Detection of poly-dA (blot hybridization) |
| Probe_2 | GGATGATCGCAGGTGAATCACCCACG | Detection of UG17 ncRNA (blot hybridization) |
| Probe_5S_E_coli | TACTCTCGCATGGGGAGACCCCAC | Detection of *E. coli* 5S RNA (blot hybridization) |
| Probe_4_6-Fam | 6-Fam-CCGAAGGTAGTAATAAGTAACGCCTG | Fluorescently labeled oligonucleotide (primer extension) |

**Table S1**. **DNA oligonucleotides used in this study.** Primer names correspond to those cited in the main text and were used for conventional PCR, inverse PCR, blot hybridization, or primer extension as indicated. Primer sequences are listed in the 5’ to 3’ orientation.
