## Supplementary material for "Host exonuclease SbcB and a phage-encoded SSB-like protein control activation of the DRT10 reverse transcriptase defense system": Table S2

| Plasmid name | Description |
|---|---|
| p57m_UG17_ncRNA-a2 | Expression of the UG17 system with a mutation in the ncRNA (GCGCCUC$_{130–136}$UGCGGAG) in the p57m vector |
| p57m_UG17_ncRNA-b2 | Expression of the UG17 system with a mutation in the ncRNA (CA$_{136–137}$UG) in the p57m vector |
| p57m_UG17-WT_RT-Flag (C-end) | Expression of the wild-type UG17 system with RT C-terminally labeled with a Flag-tag in the p57m vector |
| p57m_UG17-WT_RT-6×His (C-end) | Expression of the wild-type UG17 system with RT C-terminally labeled with a 6×His-tag in the p57m vector |
| p57m_UG17_RT-YVAA_RT-6×His (C-end) | Expression of the UG17 system with a mutation in the RT protein (DD$_{223–224}$AA) and RT C-terminally labeled with a 6×His-tag in the p57m vector |
| p57m_UG17_ncRNA-a1_RT-6×His (C-end) | Expression of the UG17 system with a mutation in the ncRNA (GAGGCGU$_{10–16}$CUCCGCA) and RT C-terminally labeled with a 6×His-tag in the p57m vector |
| p57m_UG17_ncRNA-a1&a2_RT-6×His (C-end) | Expression of the UG17 system with two mutations in the ncRNA (GAGGCGU$_{10–16}$CUCCGCA and GCGCCUC$_{130–136}$UGCGGAG) and RT C-terminally labeled with a 6×His-tag in the p57m vector |
| p57m_UG17_ncRNA-b2_RT-6×His (C-end) | Expression of the UG17 system with a mutation in the ncRNA (CA$_{136–137}$UG) and RT C-terminally labeled with a 6×His-tag in the p57m vector |
| pSparkI_UG17_ncRNA-Δ92 | Expression of the UG17 system with a 92-nt deletion in the ncRNA in the pSparkI vector |
| pBAD30_UG17_ncRNA-Δ92 | Expression of the UG17 system with a 92-nt deletion in the ncRNA in the pBAD30 vector |
| pBAD30_UG17_WT | Expression of the wild-type UG17 system in the pBAD30 vector |
| pBAD30_UG17-WT_RT-6×His (C-end) | Expression of the wild-type UG17 system with RT C-terminally labeled with a 6×His-tag in the pBAD30 vector |
| pBAD30_UG17_SLATT-Δ139_RT-6×His (C-end) | Expression of the UG17 system with a frameshift mutation in the SLATT protein and RT C-terminally labeled with a 6×His-tag in the pBAD30 vector |
| pBAD30_UG17_RT-YVAA_RT-6×His (C-end) | Expression of the UG17 system with a mutation in the RT protein (DD$_{223–224}$AA) and RT C-terminally labeled with a 6×His-tag in the pBAD30 vector |
| pBAD33_0087-WT | Expression of the wild-type *Bas52_0087* phage protein in the pBAD33 vector |
| pBAD33_0087-D252V | Expression of the *Bas52_0087* phage protein with a point mutation (D$_{252}$V) in the pBAD33 vector |
| pBAD33_0087-Y351C | Expression of the *Bas52_0087* phage protein with a point mutation (Y$_{351}$C) in the pBAD33 vector |
| pBB3_SbcB | Expression of the SbcB (ExoI) protein in the pBB3 vector |
| pBB3_SbcB15 | Expression of the SbcB15 variant with a point mutation (A$_{183}$V) in the pBB3 vector |
| pBB3_ESN2 | Expression of the wild-type ncRNA from UG17 in the pBB3 vector |

**Table S2. Plasmids constructed in this study.** Vector backbones and mutations are indicated for each plasmid.
